# Crop genotype modulates root rot resistance-associated microbial community composition and abundance of key taxa

**DOI:** 10.1101/2025.01.16.633298

**Authors:** Valentin Gfeller, Michael Schneider, Natacha Bodenhausen, Matthew W. Horton, Lukas Wille, Klaus H. Oldach, Bruno Studer, Martin Hartmann, Monika M. Messmer, Pierre Hohmann

## Abstract

Plants are constantly challenged by pathogens, which can cause substantial yield losses. The aggressiveness of and damage by pathogens depends on the host-associated microbiome, which might be shaped by plant genetics to improve resistance. How different crop genotypes modulate their microbiota when challenged by a complex of pathogens is largely unknown. Here, we investigate if and how pea (*Pisum sativum L.*) genotypes shape their root microbiota upon challenge by soil-borne pathogens and how this relates to a genotype’s resistance. Building on the phenotyping efforts of 252 pea genotypes grown in naturally infested soil, we characterized root fungi and bacteria by ITS region and 16S rRNA gene amplicon sequencing, respectively. Pea genotype markedly affected both fungal and bacterial community composition, and these genotype-specific microbiota were associated with root rot resistance. For example, genotype resistance was correlated (R^2^ = 19%) with root fungal community composition. Further, several key microbes, showing a high relative abundance, heritability, connectedness with other microbes, and correlation with plant resistance, were identified. Our findings highlight the importance of crop genotype-specific root microbiota under root rot stress and the potential of the plant to shape its associated microbiota as a second line of defense.

## 1 Introduction

Crop pathogens pose a major threat to global food security and their negative effects are expected to be further amplified by climate change (Savary et al., 2019; Singh et al., 2023). Consequently, sustainable approaches for their control are urgently needed. Microbiota-mediated pathogen resistance reflects a novel, promising tool to control crop diseases sustainably (Busby et al., 2017; Vannier et al., 2019).

Plant-associated microbiomes can promote plant health in many ways (Berendsen et al., 2012; Trivedi et al., 2020). A balanced microbiota, including bacteria, is essential for plants to protect themselves against pathogens and survive (Durán et al., 2018). To benefit from such beneficial microbiome-dependent services, plants shape the microbiota in their surrounding environment (Bulgarelli et al., 2012; Lundberg et al., 2012). This provides a promising tool to engineer a crop’s microbiome through host genetics (Escudero-Martinez and Bulgarelli, 2023). Genotype-specific plant microbiome interactions affect not only the focal plant but also the growth and defense of succeeding plants in that soil (Hu et al., 2018), which is particularly interesting in the context of crop rotation. For example, a mechanism to cope with the growth-suppressive effects of precrop-specific microbiota in the soil is the exudation of root secondary metabolites (Gfeller et al., 2024). Therefore, to enhance the resistance of grain legumes against crop rotation-specific soil-borne pathogens, the modulation of plant-microbiota interaction through breeding might also offer a promising approach (Wille et al., 2019).

Pathogen suppression by plant associated microbiota poses a promising, new approach to manage crop diseases sustainably (Busby et al., 2017; Vannier et al., 2019). It is known that the composition of the resident microbiota determines, together with the plant’s innate immune system, whether a certain microbe acts as a pathogen, commensal, or even beneficial (Entila et al., 2024; Ma et al., 2021; Pereira et al., 2023). It has been shown that plants can enrich specific taxa from the surrounding soil microbiome around their roots, thereby reducing pathogen aggressiveness and promoting plant growth (Berendsen et al., 2018; Liu et al., 2023). Banana plants, for example, can enrich beneficial fungi through specific plant exudates that confer resistance against the pathogen Foc TR4 (Liu et al., 2023). Such microbe-mediated disease resistance can be triggered by direct antagonisms like antimicrobial molecules and niche competition (Caballero-Flores et al., 2023; Chen et al., 2018; Gu et al., 2020; Matsumoto et al., 2021), or indirectly through the activation of the host immune system (Pieterse et al., 2014; Shoresh et al., 2010). Further, the endophytic root microbiota can confer induced resistance upon pathogen attack (Carrión et al., 2019). While it is established that the microbiota can mediate pathogen suppression of plants, the frequency of this phenomenon across different plants and the relative contribution of this mechanism in comparison to direct plant defense remains to be elucidated (Jian et al., 2024; Vannier et al., 2019).

Pea (*Pisum sativum,* L.) belongs to the most frequently cultivated legumes worldwide (Stagnari et al., 2017). Their symbiosis with nitrogen-fixing rhizobia can increase soil fertility, providing a benefit for themselves and subsequently cultivated crops and the peas also serve as high-quality, protein-rich food and feed (Stagnari et al., 2017; Tulbek et al., 2017). Along with other legumes, pea production is highly constrained by soil-borne pathogens, which build up upon repeated legume cropping and cause severe wilt and root rot, also known as soil fatigue, which can lead to considerable yield losses (Harveson et al., 2021; Sharma et al., 2022). Several pathogenic fungi and oomycetes can be part of the root rot complex, with the most reported causal agents being *Aphanomyces euteiches*, *Didymella pinodes*, *Didymella pinodella*, *Fusarium avenaceum*, *Fusarium oxysporum*, *Fusarium redolens*, *Fusarium solani*, *Pythium* sp., and *Rhizoctonia solani* (Alcala et al., 2016; Esmaeili Taheri et al., 2017; Gaulin et al., 2007; Harveson et al., 2021; Pflughöft et al., 2012; Sharma et al., 2022). First evidence suggests that the synergistic effect of these pathogens increases their aggressiveness, and thus, makes resistance breeding even more challenging (Baćanović-Šišić et al., 2018; Chatterton et al., 2019; Rubiales et al., 2015). Plant breeding for microbiota-dependent disease resistance could assist in overcoming this challenge (Escudero-Martinez and Bulgarelli, 2023). Screening of a pea diversity panel grown in naturally infested soil showed considerable variation in disease resistance across different pea lines (Wille et al., 2020). An in-depth analysis of eight genotypes in four different soils revealed predefined fitness-associated microbial markers as predictors for root rot resistance, including arbuscular mycorrhizal fungi (Wille et al., 2021). While microbiota communities of peas grown in healthy and infested soils have been compared and shown to differ (Chatterton et al., 2021; Hossain et al., 2021), it is largely unknown how different pea lines interact with their microbiota under root rot.

In this study, we investigate if and how pea lines with diverse resistance levels against root rot interact with their microbiota. We examined the fungal and bacterial microbiota of a diverse panel of 252 pea genotypes, consisting of gene bank accessions, breeding lines, and registered cultivars grown in a climate chamber under root rot stress. Associations of pea resistance with alpha diversity, microbial community composition, and individual taxa were investigated. Further, network and heritability analyses were performed. We found associations between microbial diversity measures and resistance levels, as well as taxa associated with resistance, which were also shown to be heritable and connected within the root microbiota. This illustrates the potential of microbiota-mediated resistance breeding against legume root rots.

## 2 Materials and Methods

### 2.1 Plant material

We capitalized on root samples collected from a previous study (Wille et al., 2020). A diverse set of 261 pea genotypes were grown under root rot stress. To eliminate confounding effects due to seed age and origin, the pea seeds were multiplied in a common environment (Sativa Rheinau AG, Switzerland) prior to the experiment. In this study, we investigated the root microbiota of a subset of 252 pea genotypes originating from three seed sources: 173 gene bank accessions of the USDA pea core collection, 33 registered European cultivars, and 46 advanced breeding lines from a Swiss plant breeder (Getreidezüchtung Peter Kunz). Two European cultivars were selected as reference lines based on their known partial resistance (EFB.33: “C1”) and susceptibility (Respect: “C2”) to root rot.

### 2.2 Characterization of root rot resistance

To study the root microbiota of the pea panel under root rot stress, a resistance assay was performed under controlled conditions as described by Wille et al. (2020). In short, soil naturally infested by pea root rot pathogens was collected on an agricultural field, and a subset was X-Ray sterilized for the control (pathogen-free) treatment. Infested and sterilized soils were mixed with autoclaved quartz sand in a 2:1 (v:v) ratio and filled into plastic pots (200 ml) to improve soil structure. Four seeds per pot were surface sterilized with ethanol and bleach and soaked in water before sowing. Each pea genotype was grown in four replicates in either infested or sterilized soil in a randomized block design. Plants were grown in a walk-in climate chamber at 20 °C, 85% relative humidity, and 16 hours of light per day. The soil was kept moist by flooding the pots every 72 hours for 30 minutes. The total number of emerged seedlings was recorded 14 days after sowing. Plants were harvested and phenotyped at 21 days. Shoot dry biomass was determined, and relative shoot dry weight was calculated by dividing the mean weight per plant in the infested soil by the mean weight in the sterilized soil (SDW_Rel_). In addition, relative total shoot biomass per pot (tSDW_Rel_) was calculated as described before but considering the biomass of all germinated plants within one pot. To evaluate belowground disease levels, plants were removed from the pots and cleaned with tap water. The root rot index (RRI: 1 = no symptoms to 6 = completely disintegrated root system) was determined, and median values per pot were used for analysis. For microbiota analysis, the previously phenotyped roots were sampled as described in Lundberg et al. (2012). Briefly, soil was removed by shaking the roots with sterile gloves, followed by a washing step in 25 ml of sterile water in a 50 ml tube by vortexing vigorously. Clean roots were stored at -20 °C until lyophilization before they were milled to fine powder in a steel jar with one 20 mm steel ball utilizing a ball mill (Retsch, Haan, Germany) at 25 Hz for 20 s.

### 2.3 Microbiota profiling

#### 2.3.1 DNA extraction and amplicon sequencing

DNA extraction was carried out with 15 mg of dry root powder using the Mag-Bind Plant DNA DS 96 Kit (Omega Bio-Tek, Norcross, United States) following the manufacturer’s instructions. The extracted DNA was diluted to 10 ng/ul and shipped to the Genome Quebec Innovation Center (Montreal, Canada) for library preparation and sequencing. For the fungal libraries, ITS1F (Gardes and Bruns, 1993) and ITS4 (White et al., 1990) primers were used to amplify the entire internal transcribed spacer (ITS) region. The amplicon libraries were prepared following the Pacific Biosciences Barcoded Universal Primers for Multiplexing Amplicons Template Preparation and Sequencing protocol. No DNA shearing was performed since the samples were amplicons, and 1’000 ng of purified amplicons were used for the library construction. The DNA Damage repair, End repair, and SMRT bell ligation steps were performed as described in the template preparation protocol with the SMRTbell Template Prep Kit 2.0 reagents (Pacific Biosciences, Menlo Park, CA, USA). The sequencing primer was annealed with sequencing primer v4 at a final concentration of 1 nM, and the Sequel II 2.1 polymerase was bound at 0.5 nM. The libraries went through an AMPure bead cleanup (following the SMRTlink calculator procedure) before being sequenced in four runs on a PacBio Sequel II instrument at a loading concentration between 120 pM and 200 pM using the diffusion loading protocol, Sequel II Sequencing kit 2.0, SMRT Cell 8M and 10 hours movies with no pre-extension. For the bacterial libraries, the V3 and V4 hypervariable regions of the 16S rRNA gene were amplified using the V3F (Chakravorty et al., 2007) and 799R (Chelius and Triplett, 2001) primers. The barcoded libraries were first normalized to 2 nM, then pooled and denatured in 0.05N NaOH. The pool was diluted to 9 pM using HT1 buffer and were loaded on a MiSeq, and sequenced in three runs for 2X300 cycles according to the manufacturer’s instructions. A phiX library was used as a control and mixed with libraries at 12% level. The MiSeq Control Software (MCS) version was 2.5.0.5, and RTA version was 1.18.54. The program bcl2fastq v1.8.4 was then used to demultiplex samples and generate fastq reads. The demultiplexed sequences have been deposited in the European Nucleotide Archive (ENA) under the accession code PRJEB83630.

#### 2.3.2 Sequence data processing

For fungal communities, the bioinformatic analysis was performed at the Genetic Diversity Centre at ETH Zurich (Switzerland). In short, raw sequences were quality-filtered and error-corrected to obtain zero radius operational taxonomic units (zOTUs) using UNOISE (Edgar, 2016a). The zOTUs were further clustered into fungal OTUs (fOTUs) of 97% nucleotide similarity with UPARSE (Edgar, 2013). Taxonomic associations were predicted using the SINTAX classifier (Edgar, 2016b) with the UNITE V8.3 (Nilsson et al., 2019) ITS reference database. For bacteria, bioinformatics was performed in QIIME2 (Bolyen et al., 2019) as previously described (Mittelstrass et al., 2021). Briefly, after importing the demultiplexed sequences, primers were removed using *cutadapt* (version 1.16 implemented in QIIME2). Trimmed reads were denoised, and exact amplicon sequences (ASVs) were inferred by utilizing the *DADA2* pipeline (Callahan et al., 2016). ASVs were then clustered into bacterial OTUs (bOTUs) of 97% nucleotide similarity using *vsearch* (v2.7.0, (Rognes et al., 2016)). Taxonomy was assigned using the SILVA database (Quast et al., 2013) and the RDP naive Bayesian classifier method implemented in DADA2 (Wang et al., 2007).

### 2.4 Statistical analysis

All statistical analyses were conducted using the open-source software R v.4.3.0 (R Core Team, 2023). Data wrangling and visualization were facilitated with the *tidyverse* package collection (Wickham et al., 2019), as well as *ggbeeswarm* and *cowplot* for visualization (Clarke et al., 2023; Wilke, 2020). The *phyloseq* R package facilitated microbiota analysis (McMurdie and Holmes, 2013). After importing bacterial and fungal count and taxonomy tables, unwanted samples or OTUs were filtered out before microbiota analysis. First, two fOTUs belonging to host DNA and the Kingdom Rhizaria and 61 bOTUs unassigned at the Kingdom level or assigned to eukaryotes, mitochondria, or chloroplasts were removed. To remove low-quality samples, five samples with less than 1’000 fungal sequences and one sample with less than 11’000 bacterial sequences were excluded. Further, 54 previously identified pea lines (Wille et al., 2020) with heterogeneous seed or flower appearance were also excluded because they were assumed to be genetically heterozygous. Lastly, low abundant OTUs were removed, if not present with at least four sequences in at least four samples. Data filtering, transformation, and analysis are summarized in **Figure S1**.

Alpha diversity was analyzed for fungi and bacteria separately by first rarefying the data to 1’000 fungal sequences and 11’000 bacterial sequences using the *vegan* package (Oksanen et al., 2020) and then calculating the OTU richness and Shannon diversity index in each sample based on the mean of 1’000 iterations. To test the effects of pea genotype and seed source (gene bank accessions, breeding material, registered cultivars) on alpha diversity, an Analysis of Variance (ANOVA) was performed. Statistical assumptions, i.e. normal distribution and homoscedasticity of error variance, were visually checked. Differences among Estimated Marginal Means (EMMs) of seed sources were tested using the *emmeans* package (Lenth, 2022), taking the Tukey method for *P* value adjustment for multiple testing and using compact letter display using the *multcomp* package for visualization (Hothorn et al., 2008). We further checked for correlations of alpha diversity with resistance-associated traits: Spearman correlations were calculated for ordinal data and Pearson correlations were performed for continuous variables.

We further examined how the microbial community composition (beta diversity) of fungal and bacterial communities relates to pea genotype, seed source, and resistance traits. Effects were visualized by Principal Coordinates Analysis (PCoA) ordination using Bray-Curtis dissimilarity matrices and tested by Permutational Multivariate Analysis of Variance (PERMANOVA) using the *adonis2* function from the *vegan* package (Oksanen et al., 2020). Multivariate homogeneity of dispersion (variance) between the seed sources was tested using the *betadisper* function from the *vegan* package (Oksanen et al., 2020). The combination of PERMANOVA and *betadisper* allows us to distinguish between location and dispersion effects in beta diversity analysis (Anderson and Walsh, 2013). We also tested whether the association between the resistance trait and the beta diversity depends on the seed source by including the interaction term of the two variables in the fitted model (model: beta diversity ∼ resistance trait * seed source). To test how the average phenotype of each genotype influences the community compositions, PCoA and PERMANVOA were also performed on mean values per genotype. Another PERMANVOA was performed to estimate the variation explained (R^2^) by the experimental replicates (blocks).

To evaluate the heritability of the root microbiota diversity indices and of the relative abundance of individual OTUs, we calculated the broad-sense heritability (H^2^). The count tables were first normalized by centered log-ratio (clr) transformation with the *aldex.clr* function implemented in the R package *ALDEx2* (Fernandes et al., 2014), as relative abundance matrices can lead to misinterpretation of OTU heritability (Bruijning et al., 2023). H^2^ was calculated as the proportion of variance explained by the random intercept effect (here, pea genotype) as described before (Brachi et al., 2022). To accomplish this, we fitted a linear mixed-effects model (y ∼ 1|’pea genotype’) using the *lme* function from the *lme4* R package (Bates et al., 2015) for alpha diversity (OTU richness and Shannon diversity index), beta diversity (PCo axis 1) and clr-transformed abundance of OTUs. To account for within-genotype variation, the replicate identity was included in the model as fixed factor. Bootstrap confidence intervals (95%) were computed using the *bootMer* function from the *lme4* R package (Bates et al., 2015) using 999 bootstraps. Differences between the median heritability of fungal and bacterial OTUs were tested by a two-sided Wilcoxon rank sum test. To assess whether there are more fungal OTUs within the 58 most heritable OTUs (H^2^ > 20%), we performed a binomial test using the *binom.test* function.

To identify OTUs that are associated with root rot resistance, we performed differential abundance analysis for individual OTUs. The R package *ALDEx2* (Fernandes et al., 2014) was chosen for the analysis as it has been shown to produce consistent results across studies (Nearing et al., 2022). Count tables were clr-transformed with the *aldex.clr* function before a Spearman correlation was performed using the *aldex.corr* function. A specific OTU was considered associated with resistance when *P* < 0.05 after Benjamini-Hochberg correction for multiple testing (Benjamini and Hochberg, 1995).

To identify highly connected (hub) OTUs, we made use of network inference. Bacterial and fungal count tables were combined before running the ‘SParse InversE Covariance Estimation for Ecological ASsociation Inference’ (SPIEC-EASI) pipeline (Kurtz et al., 2015). SPIEC-EASI was executed using the neighborhood selection method to compute the network. Network analyses were facilitated by the R package *igraph* (Csárdi et al., 2024). Betweenness centrality and degree were taken as measures for connectedness; the two measures are defined as the number of shortest paths going through a node (betweenness) and the number of its adjacent edges (degree). OTUs belonging to the top 10% of betweenness or the top 10% of degree were considered as hub OTUs.

To summarize the OTUs of most interest, we selected highly heritable (top 10%) hub OTUs that were also associated with root rot resistance (top 10% in correlations with at least one resistance-associated trait) and showed a relative abundance above 1%. These OTUs were further taxonomically investigated. Besides the above-mentioned taxonomic assignment, we further investigated the taxonomy by performing a standard nucleotide blast to the NCBI database. If more than one taxonomic entry showed the maximum percentage identity at the maximum query coverage, the next higher taxonomic level was indicated. The same procedure was performed for OTUs belonging to taxonomic groups that are expected to be involved in root rot and resistance to it (Wille et al., 2021).

We further investigated how best to model disease resistance using measures of OTU abundance and beta diversity. Seedling emergence and RRI were modeled through four models: (i) A stepwise regression on the 20 most abundant OTUs and the two first PCo axes (beta diversity) of both domains (bacteria and fungi), performed with the *stepAIC* function from the *MASS* package (Ripley et al., 2024) using backward selection direction; (ii) A stepwise regression on these (40) OTUs only; (iii) A simple linear model with the PCo axis showing the highest correlation; (iv) A simple linear model with the OTU showing the highest correlation. The models were compared based on Akaike’s information criterion (AIC), adjusted R^2^ and the corresponding *P* value.

## 3 Results

### 3.1 Microbial diversity is affected by pea genotype under root rot stress

Amplicon sequencing provided 6’726 ± 4’294 (mean± SD) fungal and 26’471 ± 6’781 bacterial sequences per sample, with variable coverage across sequencing runs calling for data normalization prior to downstream analysis (**Figure S1**, **Figure S2**). A first inspection of the most abundant taxa in our data revealed that the most abundant phyla were Ascomycota in the fungal Kingdom and Proteobacteria in the bacterial domain (**Figure S3**). We found a few genera that contributed to a large fraction of the microbiota. For Fungi, 19% of the reads were assigned to the genus *Dactylonectria* and 12% to *Fusarium*. For Bacteria, 49% of the reads were assigned to possibly nodule-inhabiting *Rhizobium* spp. (SILVA databank genus taxonomy: “Allorhizobium-Neorhizobium-Pararhizobium-Rhizobium”), 15% to *Flavobacterium* spp. and 9% to *Pseudomonas* spp.

To examine the variation within our data set, we first compared alpha diversity indices among pea genotypes and seed sources, where the seed source reflects different levels of breeding intensity. The plant genotype explained 16% and 25% of the variation in fungal, respectively bacterial Shannon diversity, whereas the seed source explained only 1-2% of the variation (**Figure 1**). Similar patterns were observed for OTU richness (**Figure S4)**. The most consistent difference in alpha diversity among the seed sources was found between the gene bank material and the breeding material, where Shannon diversity and OTU richness were reduced in the genotypes originating from the gene bank.

**Figure 1.**
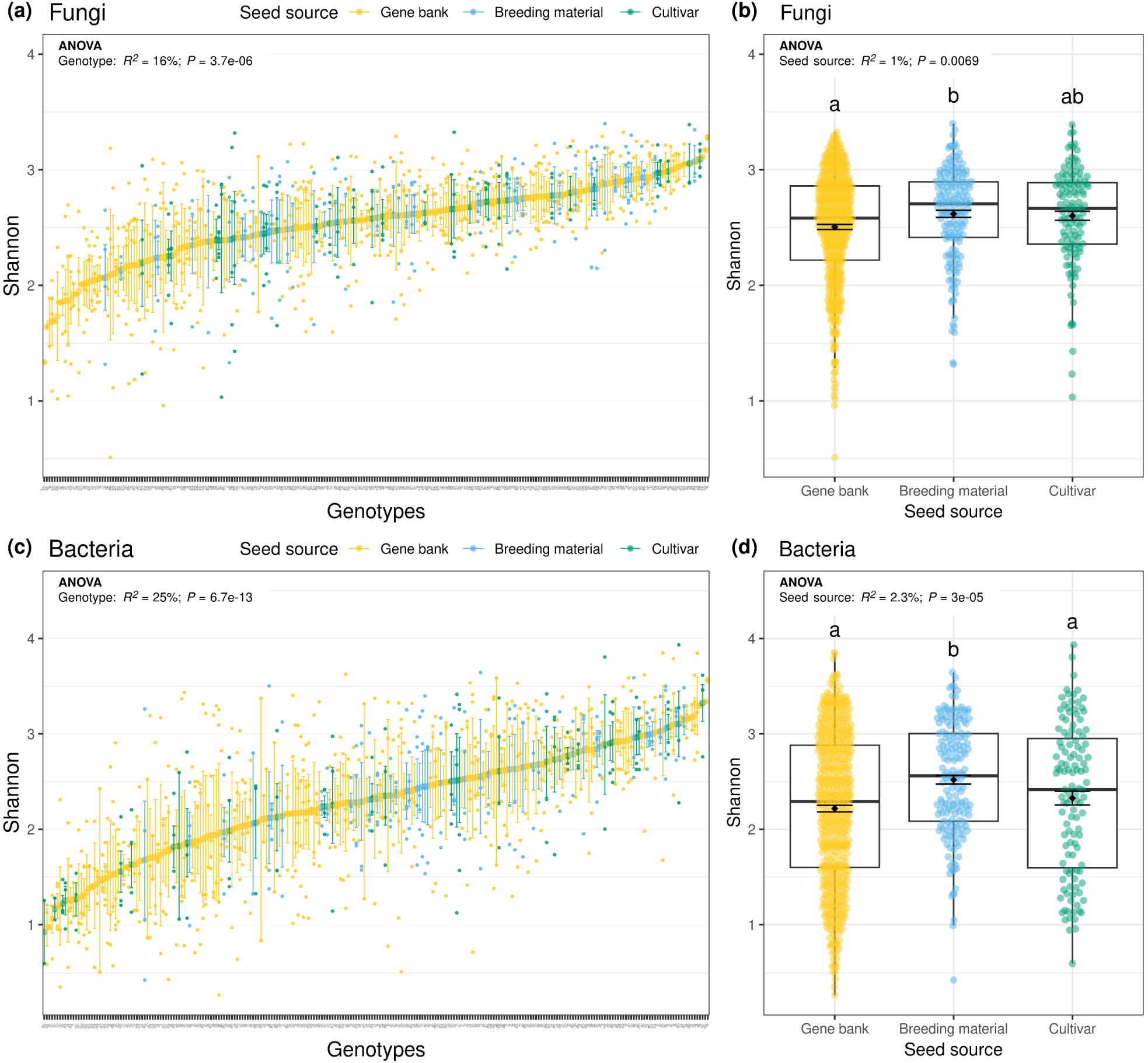
Influence of plant genotype **(a, c)** and seed source **(b, d)** on root fungal **(a, b)** and bacterial **(c, d)** alpha diversity. All plots show individual datapoints, means ±SE, and the ANOVA results with the explained variance (adjusted R^2^) and the corresponding *P* value. In **(b)** and **(d)** boxplots are also shown. Letters indicate significant differences among seed sources (analysis of variance followed by pairwise comparison of estimated marginal means, *P*_adj_ < 0.05).

Similar to alpha diversity, the plant genotype explained a large fraction of the microbial beta diversity (45-51%), whereas the influence of the seed source was relatively small (1-4%) (**Figure 2**), as revealed by PERMANOVA. Visualization of the first two axes of a Principal Coordinates Analysis (PCoA) confirmed the spatial separation of the two reference genotypes (EFB.33 and Respect) and the absence of a strong seed source effect. For fungi, gene bank accessions also showed an enhanced multivariate dispersion compared to the other seed sources (**Figure S5**). For unbalanced designs where the larger group has a greater dispersion, as in this case, the test might be overly conservative (Anderson and Walsh, 2013).

**Figure 2.**
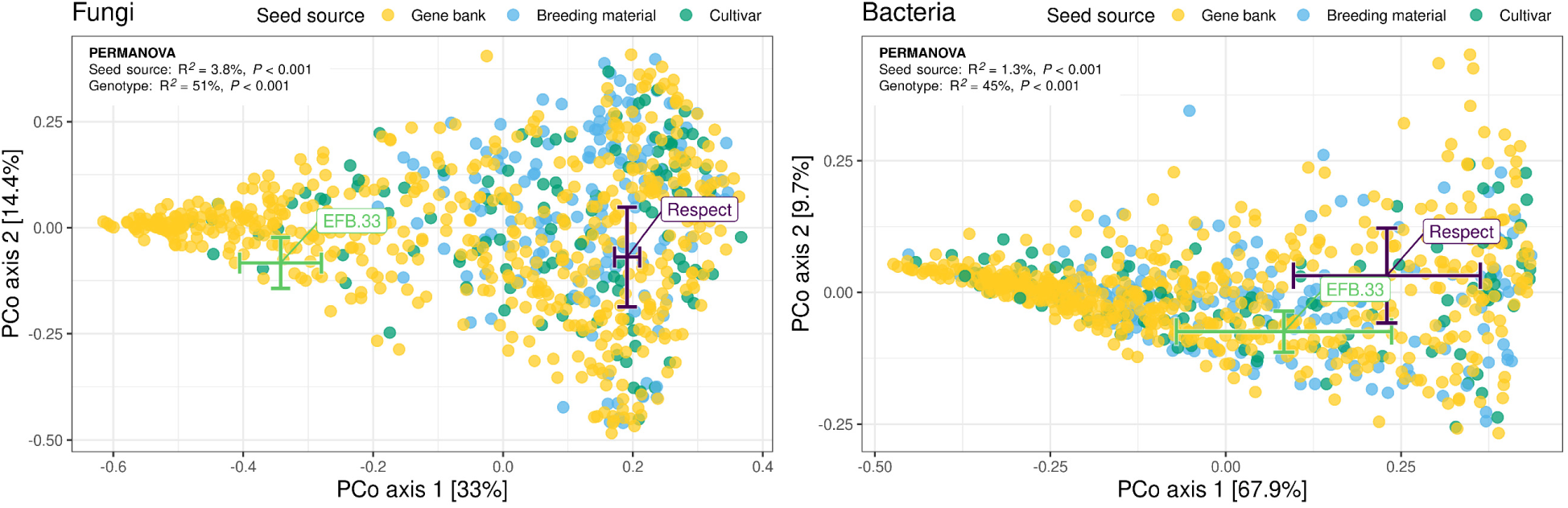
Influence of plant genotype and seed source on root microbial beta diversity assessed by Principal Coordinates Analysis (PCoA) ordination of fungal and bacterial communities. Individual datapoints and means ±SE for two reference pea genotypes (EFB.33, Respect) are shown. PERMANOVA results with the explained variance (R^2^) and the corresponding *P* value are included in both plots.

### 3.2 Variation in microbial diversity is associated with root rot resistance

To evaluate the interplay between disease resistance and root microbial diversity, we tested for associations between alpha diversity indices and root rot phenotypes (emergence and RRI). For bacteria, an increase in Shannon diversity and OTU richness was associated with reduced seedling emergence and higher RRI, which was most pronounced for Shannon diversity and RRI (R^2^ = 9.4%, **Figure S6, Figure S7**). In contrast, less striking and inconsistent associations were found for fungi.

PERMANOVA and visualization of PCoA revealed a significant link between all resistant-associated traits and beta diversity in both domains (**Figure 3a,b**). For fungi, traits with the strongest association were found to be emergence (R^2^ = 9.3%) and relative total shoot dry weight per pot (tSDW_Rel_, R^2^ = 9.4%; **Figure 3a**). In contrast, for bacterial communities, the strongest association was found for RRI (R^2^ = 8.6%, **Figure 3b**). When calculated on mean values per genotype, the explained variation for microbial community composition increased approximately two-fold for fungi (R^2^_emergence_ = 19%, **Figure 3c**) and bacteria (R^2^_RRI_ = 14%, **Figure S8**), demonstrating that there was considerable within-genotype variation (replicate and stochastic effects) in our study system. Another PERMANVOA revealed that the experimental replicate only showed a minor but statistically significant effect on the community composition (fungi: R^2^_replicate_ = 0.97%; bacteria: R^2^_replicate_ = 1.9%). Interestingly, we found that the association between emergence and fungal community composition depended on the seed source, as shown by a statistically significant interaction term between the two variables in PERMANOVA (R^2^_emergence x ‘seed source’_ = 2.2%, **Figure 3a,c**). No such dependence on seed source was found for bacterial communities (**Figure S8**). To further disentangle the interdependency of resistance and seed source in fungal beta diversity, we performed an ANOVA followed by individual correlation analyses for each seed source between community composition (PCo axis 1) and emergence (**Figure 3d**). Again, we observed that beta diversity is associated with seed source, emergence, and their interaction. Correlating community composition and emergence within each seed source separately, we found a statistically significant link for the gene bank material (R^2^ = 3.6%) and cultivars (R^2^ = 3.4%) but not for the breeding population.

**Figure 3.**
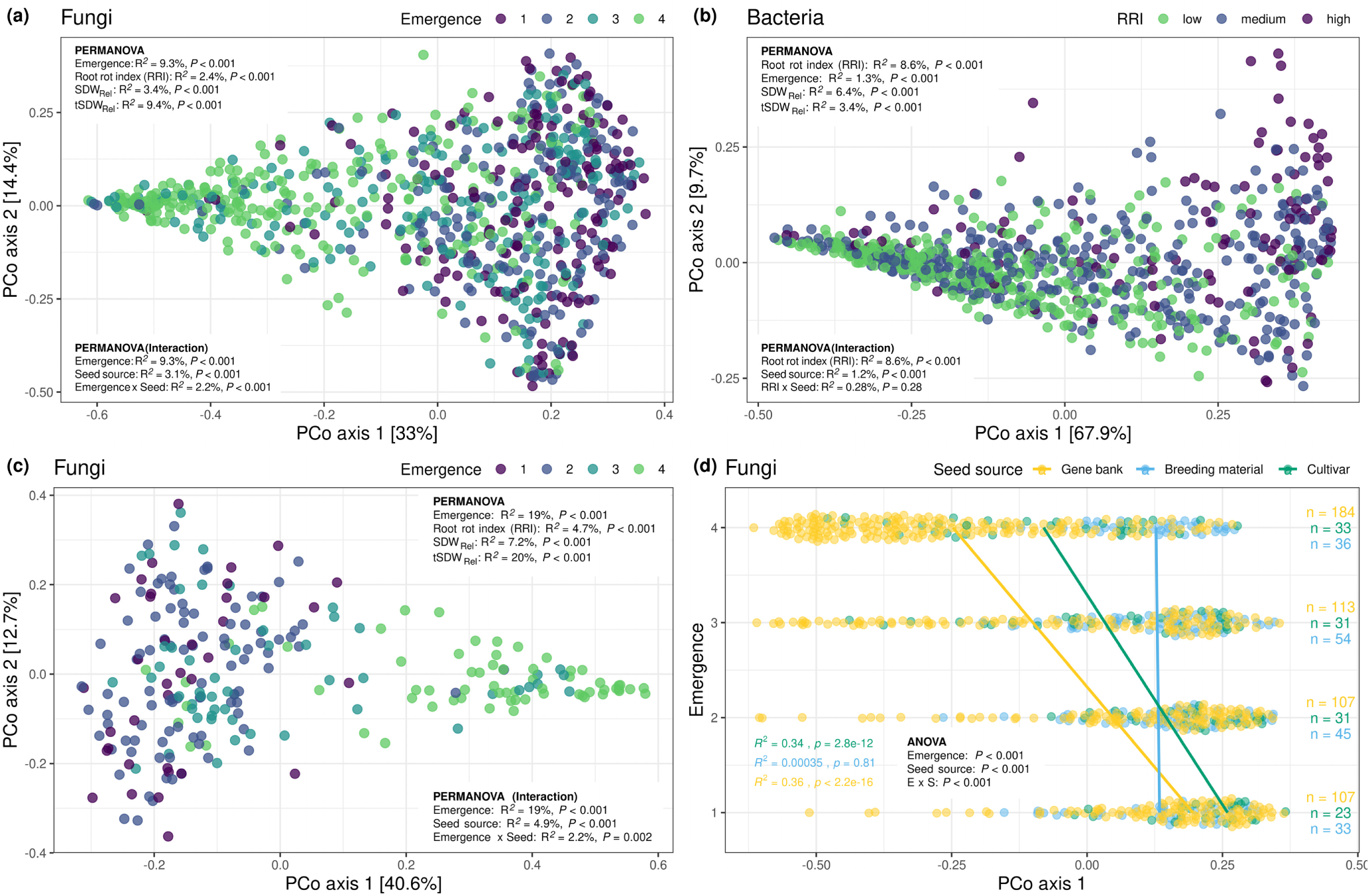
Association of root fungal **(a**, **c**, **d)** and bacterial **(b)** beta diversity with root rot resistance assessed by Principal coordinates Analysis (PCoA) ordination. **(a, b, c)** PERMANOVA results with the explained variances (R^2^) and the corresponding *P* values for different root rot resistance-associated traits (top) and the interaction with seed source (bottom) re included. **(c)** PCoA and PERMANOVA results on means per genotype. **(d)** Correlation of emergence and fungal PCo axis for each seed source. ANOVA table, as well as R^2^ and *P* values of correlations for all seed sources are shown. The number of samples per emergence level and seed source are also indicated (n). SDW_Rel_: Relative shoot dry weight per plant, tSDW_Rel_: elative total shoot dry weight per pot.

### 3.3 Microbial diversity and resistance-associated OTUs are heritable

For alpha diversity indices (Shannon and OTU richness), the broad-sense heritability (H^2^) varied between 11% and 25%, with similar levels in both domains (**Figure 4a**). By contrast, H^2^ of beta diversity, summarized by PCo axis 1 and 2, was highly domain-specific (**Figure 4a**). Whereas H^2^ of bacterial community composition (R^2^ = 25.5%) was comparable to alpha diversity heritability, H^2^ of the fungal PCo axis 1 was strikingly high (R^2^ = 70%).

**Figure 4.**
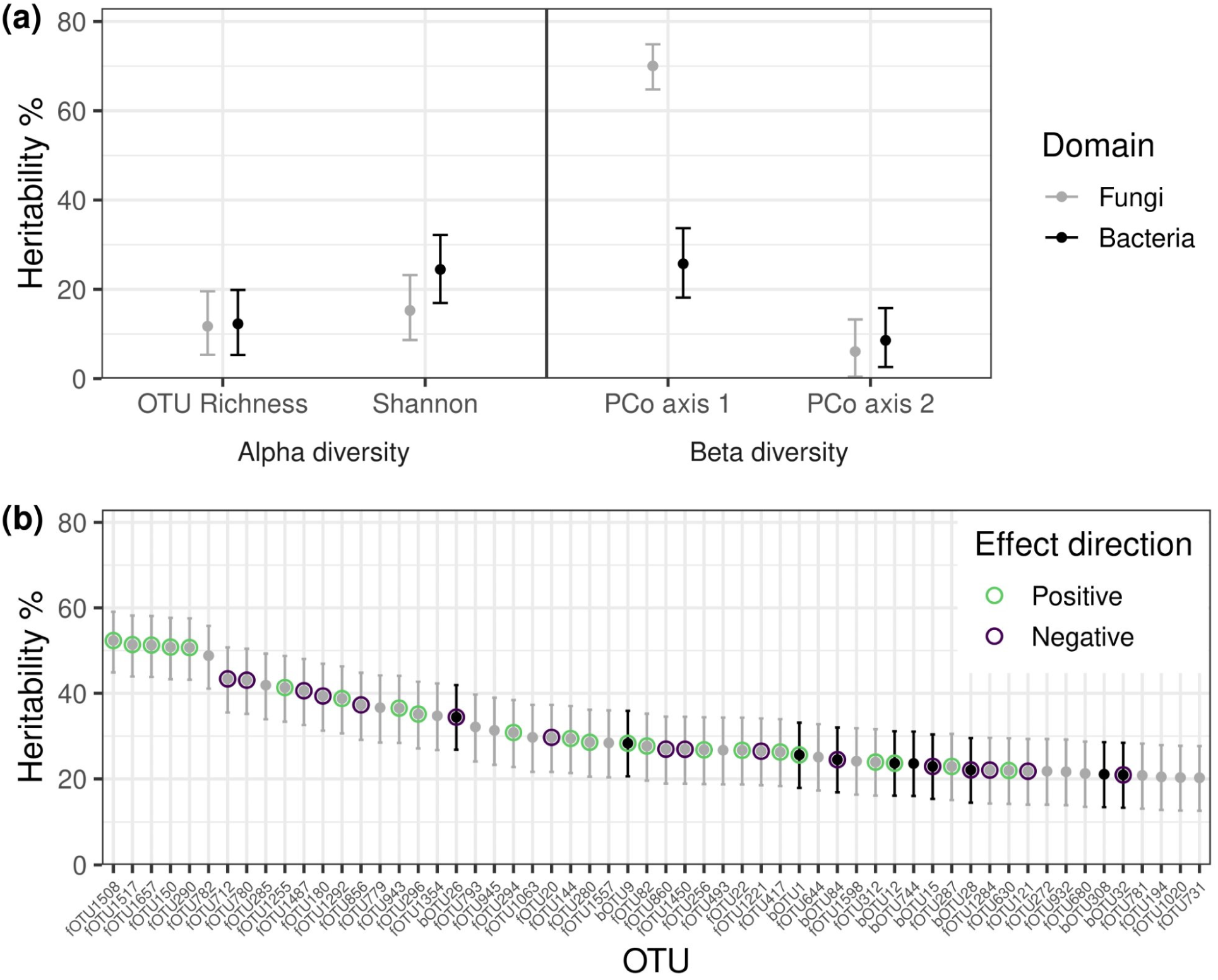
Broad-sense heritability (H^2^) of root microbial diversity and individual OTUs. **(a)** H^2^ of alpha and beta diversity of fungi and bacteria. **(b)** H^2^ of the 58 top heritable OTUs (H^2^ > 20%), including fungi (grey) and bacteria (black). OTUs positively (green) or negatively (purple) correlated with disease resistance are highlighted. **(a, b)** Bootstrap confidence intervals (95%) for H^2^ are shown.

Fifty-eight OTUs showed an H^2^ above 20% (**Figure 4b**). Overall, fungal OTUs were more heritable (median H^2^ = 4.62%) than bacterial OTUs (median H^2^ = 2.3%; *P* < 0.001). Also, among the most heritable OTUs, there were significantly more fungi than bacteria; 48 out of the top 58 OTUs were fungi (*P* < 0.001). With a H^2^ of 52%, fOTU1508 showed the highest H^2^.

Fungal and bacterial OTUs were screened for their association with root rot resistance using the *ALDEx2* differential abundance pipeline. Fifty-one fungal and 34 bacterial OTUs were found to be associated with all resistance traits (**Figure 5a,b**). These core differential abundant OTUs either correlated with plant resistance (fungi: 35, bacteria: 13) or plant susceptibility (fungi: 17, bacteria: 21; **Figure 5c,d**). Similarly to beta diversity, the highest number of resistance-associated OTUs were found for the traits emergence and tSDW_Rel_ for fungi and RRI for bacteria (**Figure 5a,b**; **Figure 3**). Identical to the heritability analysis, fOTU1508 also exhibited the strongest association with root rot resistance (R^2^ = 24.5%).

**Figure 5.**
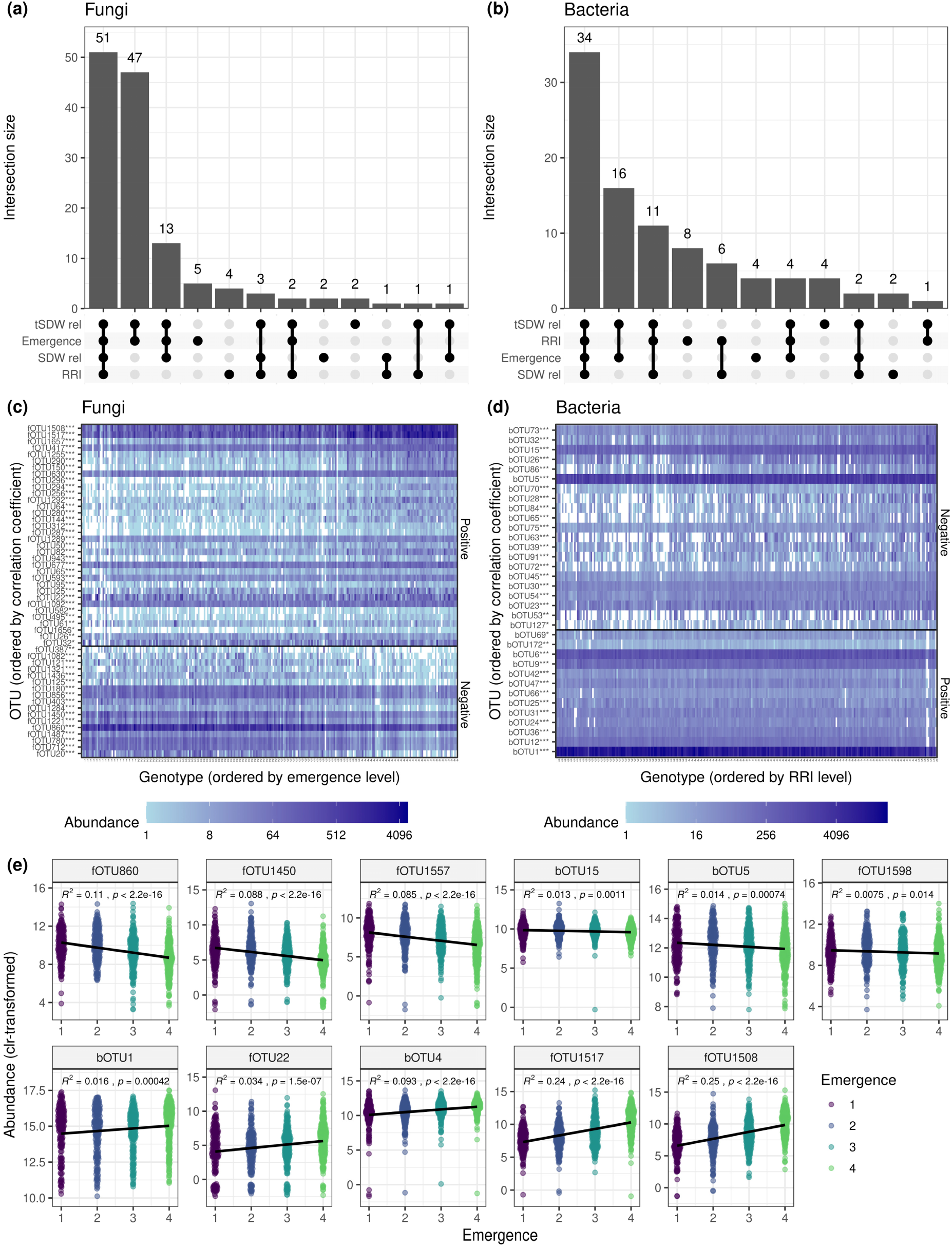
Association of fungal **(a, c, e)** and bacterial **(b, d, e)** OTUs with root rot resistance traits. **(a, b)**. UpSet plots show the number of OTUs that are significantly correlated to root rot-associated traits and their intersection, where the first bar shows the OTUs that are significant for all traits. **(c, d)** Heats map indicating the abundance OTUs significant for all traits, where the significance level is indicated with asterisks. The solid black line represents the transition from positive to negative correlation. Genotypes are ordered by increasing **(c)** emergence and **(d)** increasing RRI, and OTUs are ordered by their strength of correlation with the resistance trait. **(e)** Correlation of emergence and OTU abundance for OTUs with high relative abundance, connectedness in the microbial network, heritability and association with resistance. Explained variance (R^2^) and *P* values of correlations are shown. SDW_Rel_: Relative shoot dry weight per plant, tSDW_Rel_: Relative total shoot dry weight per pot, RRI: Root rot index. Levels of significance: *P* < 0.001 ***, *P* < 0.01 **, *P* < 0.05 *.

Given the overlap of highly heritable OTUs and root rot-associated OTUs, we tested the hypothesis that H^2^ is correlated with the magnitude of root rot resistance-association. Indeed, we found that an increase in root rot resistance-association was correlated with an increase in H^2^ (**Figure S9**). This was observed to a similar extent for resistance-(R^2^ = 68%) and susceptibility-associated OTUs (R^2^ = 69%).

Network inference using the *SPIEC-EASI* pipeline revealed little overlap between nodes (OTUs) of the two domains (**Figure S10**). Further, most of the highly heritable and differential abundant nodes are co-locate within the network. Considering OTUs belonging to the top 10% (122 OTUs) of either betweenness or degree we identified 172 hub OTUs.

To identify OTUs of potential high relevance, we selected hub OTUs that were (i) highly heritable (top 10%), (ii) highly linked to root rot resistance (top 10% of correlations of at least one resistance-associated trait), and (iii) abundant (relative abundance > 1%). Based on these requirements, we found 11 OTUs, seven fungi, and four bacteria (**Table 1**). Interestingly, all of these OTUs also showed a high prevalence, indicating that they were detected in most of the samples (ranging from 91.2% to 100%), while the majority of other OTUs were found to be less prevalent (**Figure S11**). Visualization of the association between OTU abundance and disease resistance of the selected OTUs further highlights that fungi are more associated (higher R^2^) with emergence, whereas bacteria are more associated (higher R^2^) with RRI (**Figure 5e** and **Figure S12**).

**Table 1.** Attributes of most root rot associated, heritable, and interconnected OTUs. Taxonomy, relative abundance (RA), heritability, prevalence, network characteristics (degree and s), and correlation (Spearman) with root rot resistance-associated traits of OTUs are listed. *P* values < 0.05 are highlighted in bold.

| OTU id | Phylum | Family | Genus | Blast hit<sup>A</sup> | RA (%)<sup>B</sup> | Heritability (%) | Prevalence (%) | Degree | Betweenness | Emergence |  | RRI |  |
| --- | --- | --- | --- | --- | --- | --- | --- | --- | --- | --- | --- | --- | --- |
|  |  |  |  |  |  |  |  |  |  | cor | *P* | cor | *P* |
| fOTU860 | Ascomycota | Nectriaceae | Fusarium | Fusarium sp. | 10.8 | 26.9 | 100 | 21 | 5950 | -0.322 | < **0.001** | 0.118 | **0.006** |
| fOTU1450 | Ascomycota | Nectriaceae | unassigned | Fusarium sp. | 1.33 | 26.8 | 96.7 | 21 | 3980 | -0.291 | < **0.001** | 0.168 | < **0.001** |
| fOTU1557 | Ascomycota | unassigned | unassigned | Helotiales (Order) | 2.64 | 28.3 | 99 | 19 | 6590 | -0.288 | < **0.001** | -0.06 | 0.215 |
| bOTU15 | Proteobacteria | Rhizobiaceae | Rhizobium* | Rhizobium sp. | 1.47 | 23 | 99.9 | 67 | 17900 | -0.124 | **0.006** | 0.302 | < **0.001** |
| bOTU5 | Proteobacteria | Pseudomonadaceae | Pseudomonas | Pseudomonas sp. | 8.96 | 18.8 | 100 | 36 | 10000 | -0.123 | **0.007** | 0.244 | < **0.001** |
| fOTU1598 | unassigned | unassigned | unassigned | Fusarium sp. | 8.84 | 24.2 | 100 | 22 | 5600 | -0.086 | **0.043** | -0.118 | **0.006** |
| bOTU1 | Proteobacteria | Rhizobiaceae | Rhizobium* | Rhizobium sp. | 45.7 | 25.6 | 100 | 44 | 14100 | 0.139 | **0.001** | -0.35 | < **0.001** |
| fOTU22 | unassigned | unassigned | unassigned | Olpidium sp. | 1.46 | 26.5 | 91.1 | 20 | 4290 | 0.182 | < **0.001** | -0.35 | < **0.001** |
| bOTU4 | Actinobacteriota | Streptomycetaceae | Streptomyces | Streptomyces sp. | 3.46 | 13.7 | 99.3 | 34 | 7100 | 0.295 | < **0.001** | -0.052 | 0.471 |
| fOTU1517 | Ascomycota | Nectriaceae | Dactylonectria | Dactylonectria sp. | 11.7 | 51.1 | 99.5 | 25 | 6660 | 0.488 | < **0.001** | -0.267 | < **0.001** |
| fOTU1508 | Ascomycota | Nectriaceae | Dactylonectria | Dactylonectria sp. | 8.31 | 52.3 | 99.5 | 36 | 8970 | 0.495 | < **0.001** | -0.322 | < **0.001** |

<sup>A</sup>Top hit of a BLAST search against the NCBI database; <sup>B</sup>Within-Kingdom relative abundance was computed for each OTU;
\*`Rhizobium` refers to the genus `Allorhizobium-Neorhizobium-Pararhizobium-Rhizobium`

### 3.4 Resistance- and susceptibility-associated OTUs are taxonomically diverse

To get a better understanding of the identified relevant resistance- and susceptibility-associated OTUs, we further investigated their taxonomic assignment. For fungi, the plant resistance-associated OTUs were *Dactylonectria* spp. (fOTU1508, fOTU1517; r = 0.5 and 0.49, *P* < 0.001) and *Olpidium* sp. (fOTU22; r = 0.18, *P* < 0.001; **Table 1**), the disease-associated OTUs were two *Fusarium* spp. (fOTU860, fOTU1450; r = -0.32 and -0.29, *P* < 0.001) and an OTU belonging to the Order Helotiales (fOTU1557; r = -0.29, *P* < 0.001). One putative *Fusarium* sp. (fOTU1598; r = -0.09, *P* = 0.043) was negatively correlated with emergence and RRI, thus associated with susceptibility (emergence) and resistance (RRI). For bacteria, the plant resistance-associated OTUs were a *Streptomyces* sp. (bOTU4; r = 0.3, *P* < 0.001) and a highly abundant, therefore likely nodule-inhabiting, *Rhizobium* sp. (bOTU1; r = 0.14, *P* = 0.001), and the disease-associated OTUs were another *Rhizobium* sp. (bOTU15; r = -0.12, *P* = 0.006) and a *Pseudomonas* sp. (bOTU5; r = -0.12, *P* = 0.007).

We further identified OTUs within our dataset belonging to taxa that are expected to relate to root rot and resistance to it (Wille et al., 2021). Several of these additional candidate taxa were significantly associated with emergence and/or RRI (**Table S1**). As potential beneficial microbes, we searched for OTUs related to Arbuscular mycorrhizal fungi (AMF) and *Clonostachys rosea*. We found one OTU assigned to a *Funneliformis* sp. (fOTU1020, AMF), which was negatively correlated with RRI (r = -0.24, *P* < 0.001), and one putative *Clonostachys rosea* (fOTU60), which was weakly correlated with higher emergence (r = 0.1, *P* = 0.018; **Table S1**). As potential members of the root rot complex, we identified six *Fusarium* spp. (fOTU419, fOTU680, fOTU329, fOTU334, fOTU137, fOTU487) besides the ones reported above, one putative *Didymella pisi* (fOTU40; teleomorph of *Ascochyta pisi*), and four OTUs assigned to *Rhizoctonia* (fOTU831, fOTU578, fOTU140) or the synonymous genus *Ceratobasidium* (fOTU335; Oberwinkler et al., 2013). One *Fusarium* sp. (fOTU419; r = -0.14, *P* <0.001), one *Rhizoctonia* sp. (fOTU831; r = -0.14, *P* = 0.006), and the putative *Ascochyta pisi* (fOTU40; r = -0.15, *P* < 0.001) were more abundant in low-emerging samples. Another *Fusarium* sp. (fOTU137, putative *Fusarium redolens*; r = -0.09, *P* = 0.05) was negatively correlated with RRI, and the remaining seven identified OTUs did not show any significant correlation.

### 3.5 Improved modeling of plant resistance through a combination of multiple OTUs and beta diversity

Comparing univariate models and stepwise regression models with OTU abundance and beta diversity revealed that 15 and 19 variables best explained (low AIC, high R^2^) the disease resistance phenotypes emergence and RRI, respectively (**Table S2, Table S3**). For emergence, the explained variance of the univariate models with the most correlated OTU (fOTU1508, R^2^ = 22.8%) or the most correlated PCo axis (fungal PCo axis 1, R^2^ = 27.5%) performed slightly worse compared to the stepwise regression OTU models without (R^2^ = 32.7%) and with beta diversity (R^2^ = 36.7%). For the RRI, the explained variance of the univariate models was markedly lower (bOTU1, R^2^ = 15%; bacterial PCo axis 1, R^2^ = 10.6%), while the stepwise regression model explained 33.9% (R^2^) of the variance. Adding beta diversity measures did not increase model performance for RRI (**Table S3**).

## 4 Discussion

Plants are known to shape their associated microbiota in a genotype-specific manner (Bulgarelli et al., 2012; Horton et al., 2014), which in turn affect plant growth and defense (Berendsen et al., 2012; Pieterse et al., 2014). To what extent microbiota attributes are associated with complex plant traits such as root rot resistance is largely unknown. The resistance of a given plant genotype against root rot was proposed to be influenced by individual known beneficial microbes (Wille et al., 2021, 2019). However, the extent to which the wider microbial community is associated with a genotype resistance is poorly understood. Here, we analyzed the root bacterial and fungal microbiota of a diverse set of pea genotypes grown under root rot stress and demonstrated that, in addition to individual key taxa, community-wide microbiota attributes are also associated with root rot resistance.

Our study reveals that associations, e.g. between root rot resistance and fungal and bacterial community composition, are driven by the individual pea genotypes. In line with our findings, common bean cultivars resistant to an individual fungal root pathogen (*Fusarium oxysporum)* were shown to harbor a distinct microbiota composition that is different from susceptible cultivars (Mendes et al., 2018). Our results expand these findings by investigating more than 250 genotypes in a field soil naturally infested with several pathogens, allowing us to show the range of microbial communities shaped by pea plants grown in the same soil. Notably, associations of fungal but not bacterial beta diversity were dependent on the seed source, with no correlation found for the Swiss breeding material (**Figure 3**). This seed source dependency highlights the importance of screening plant microbiota interactions of large diversity panels. We demonstrate that the seed source effects are neglectable (alpha: 1-2.3%, beta: 1.3-3.8%) relative to the genotype effect (alpha diversity R^2^: 16-25%, beta diversity R^2^: 45-51%), even though within our panel the genotypic structure and phenotypic appearance of the modern European cultivars differ substantially from the gene bank accessions that mostly consists of landraces from around the world (Ariza-Suarez et al., 2024; Wille et al., 2020). Compared to the literature, the proportion of variance in beta diversity explained by the plant genotype was exceptionally high in our study (Wagner, 2021); in diversity panels of sorghum and maize, for example, the genotype effect on root microbiota composition was found to be 7.5% and 9.8% (Deng et al., 2021; He et al., 2024). This could be due to stress conditions increasing the heritability of the root-associated microbiota (He et al., 2024). Further, we report an increase in multivariate dispersion for gene bank accessions, indicative of a more variable set of microbiota communities. Thus, gene bank accessions could pose an interesting reservoir for microbiome-recruiting/manipulation alleles that are not present in breeding material or varieties, thereby supporting the breeding of microbiome-smart cultivars for sustainable agriculture (Nerva et al., 2022). However, given that the analyzed cultivars cover most of the microbial variation, they could also be used as a first source of desired alleles to speed up the introgression into elite cultivars due to the reduced number of unwanted alleles compared to gene bank accessions.

The abundance of individual OTUs can be determined by host genotype and have an impact on plant fitness (Brachi et al., 2022). In this study, we report multiple fungal and to some extent also bacterial OTUs that are associated with disease resistance or susceptibility (**Figure 5)**. The biggest portion of them is correlated with all the measured resistance phenotypes (root rot index (RRI), emergence, relative shoot dry weight per plant/pot), reflecting the magnitude of pathogen damage (RRI) as well as several plant performance measures. Given that root rot in peas is expected to be caused by fungal and oomycetous pathogens (Harveson et al., 2021), the many bacterial OTUs found to be positively associated with RRI might have passively entered the roots through lesions or assisted as pathogen helper bacteria (Li et al., 2022). The higher bacterial alpha diversity in roots of diseased plants supports the hypothesis that bacteria can enter diseased roots more easily (**Figure S6, Figure S7**). In contrast, the resistance-associated bacterial and also fungal OTUs might directly or indirectly promote the plant health status or merely be spurious. Irrespective of a causal link, these key taxa show promise as resistance indicators in selection assays. It is striking that the OTUs that are correlated the most with resistance share several attributes: they generally have a high relative abundance, are highly interlinked with other OTUs (hub taxa), and are highly heritable (**Table 1**). The heritability of individual OTUs was markedly correlated with the degree of positive or negative disease association (**Fig. S9**), which could be explained either by root rot severity-dependent microbial root colonization (Larousse et al., 2017) or by microbiota-mediated resistance to root pathogens (Jian et al., 2024; Vannier et al., 2019).

Investigating the taxonomy of the key OTUs (**Table 1**) revealed that the two most resistance-associated OTUs (fOTU1508, fOTU1517) belong to the genus of *Dactylonectria*. Various members of this genus have been associated with severe root disease in many annual and perennial plants like strawberries and grapevines (CARLUCCI et al., 2017; Manici et al., 2018; Weber and Entrop, 2017). Other studies found *Dactylonectria* spp. in asymptomatic roots of different plant species (Berlanas et al., 2020; Durán et al., 2018), including the legume soybean, where members of this genus were shown to be enriched in the root endophytic community (Moroenyane et al., 2021; Strom et al., 2020). To the best of our knowledge, this is the first report of *Dactylonectria* spp. being associated with disease resistance, highlighting this and other identified taxa as potential microbial markers to screen for root rot resistance in peas. The most susceptibility-associated OTUs were found to be *Fusarium* species. This underlines the well-investigated role of *Fusarium* spp. (e.g. *F. solani*, *F. oxysporum*) among the most prominent pathogens that cause pea root rot worldwide (Bani et al., 2018; Harveson et al., 2021). We also identified several additional taxa associated with root rot, suggesting that an even broader range of microbes may play a role in triggering root rot. Future experiments in different soils and under field conditions will show if the predictive power of the identified microbial markers can be translated to other environments. This will further evaluate the potential of microbiota-assisted disease resistance to improve agroecosystem sustainability.

Plant resistance is well-known to be heritable (Hammond-Kosack and Jones, 1996). In our experimental system, the resistance to pea root pathogens was found to be highly heritable for emergence (H^2^ = 89%), followed by relative shoot dry weight (H^2^ = 51%) and root rot index (H^2^ = 43%, Wille et al., 2020). For the root microbiota, we report similar heritability levels of up to 52% for individual resistance-associated OTUs and 70% for the fungal community composition (beta diversity; **Figure 4**). The abundance of single OTUs explained up to 22.8% (adjusted R^2^) of the disease severity (**Table S2**). Combining the abundance of several OTUs with microbial community composition (PCoA axes) improved the prediction of root rot resistance to 36.7%. Incorporating information about the microbiota as decision support in the selection process of plant breeding could assist in promoting the beneficial root microbiota as a second line of defense against root pathogens. Genome-wide association studies and genomic prediction analysis will enable the identification of genetic loci associated with the potential recruitment of beneficial microbes and evaluate the joint use of microbial and plant markers (Escudero-Martinez and Bulgarelli, 2023). Together, this will further pave the way for microbiome-assisted breeding to overcome the challenges currently faced in resistance breeding against root rot in legumes.

## Acknowledgements

We thank the Genetic Diversity Centre (GDC) at ETH Zurich for their contribution to the processing of the raw sequencing reads. This research was supported by the Gebert Rüf Foundation (GRS-082/19), Root2Res (EU Horizon Europe no. 101060124 and Swiss State Secretariat for Education, Research and Innovation (SERI) no 23.00050), LIVESEED (EU Horizon 2020 no. 727230 and SERI no. 17.0009) and MICIU/AEI/10.13039/501100011033 and FSE+ under grant reference no RYC2022-037997-I. The opinions expressed and arguments employed herein do not necessarily reflect the official views of the EC and the Swiss government. Neither the European Commission, SERI nor any person acting on behalf of the Commission/SERI is responsible for the use which might be made of the information provided.

## Data availability statements

The raw sequencing data can be downloaded from the European Nucleotide Archive (ENA, http://www.ebi.ac.uk/ena) under the study accession PRJEB83630. R code and data to reproduce the statistical analysis and visualizations are available under https://github.com/ValentinGfeller/Gfeller_et_al_Microbiota_pea_panel.

## Author contributions

MM, PH, and BS designed research. LW performed research. VG and MWH (QIIME2) analyzed data with input from MH, NB, MS, and PH. VG wrote the first draft of the manuscripts with input from PH, MH and MM. MM, MH, NB, MS, PH, KO contributed to the interpretation of the results and the writing of the manuscript. All authors approved the final version of the paper.

## Supporting Information

### Supplementary Figures

**Figure S1.**
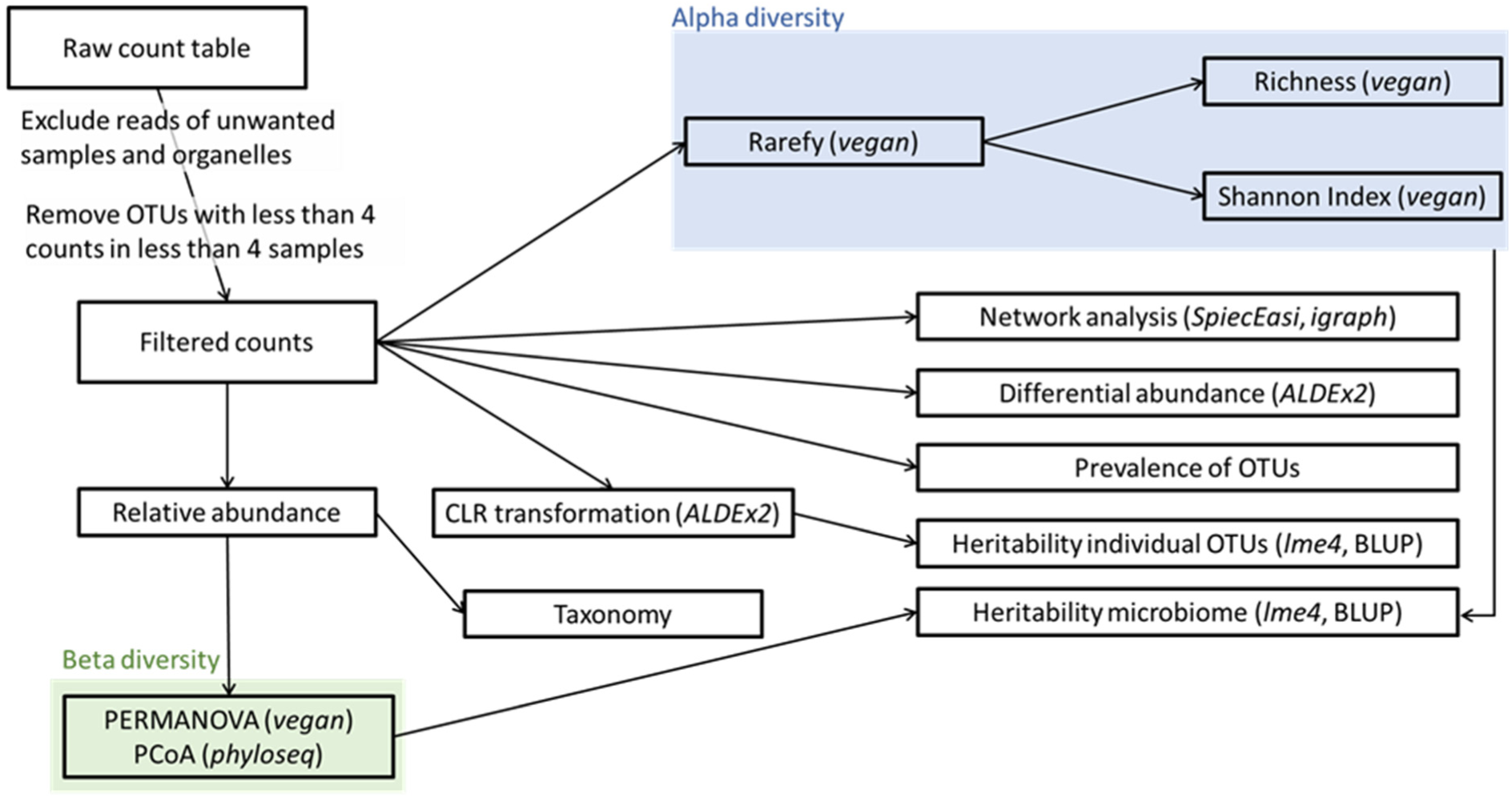
Overview of data filtering, transformation and analysis. R package names are indicated in italics. Details can be found in the Materials and Methods section.

**Figure S2.**
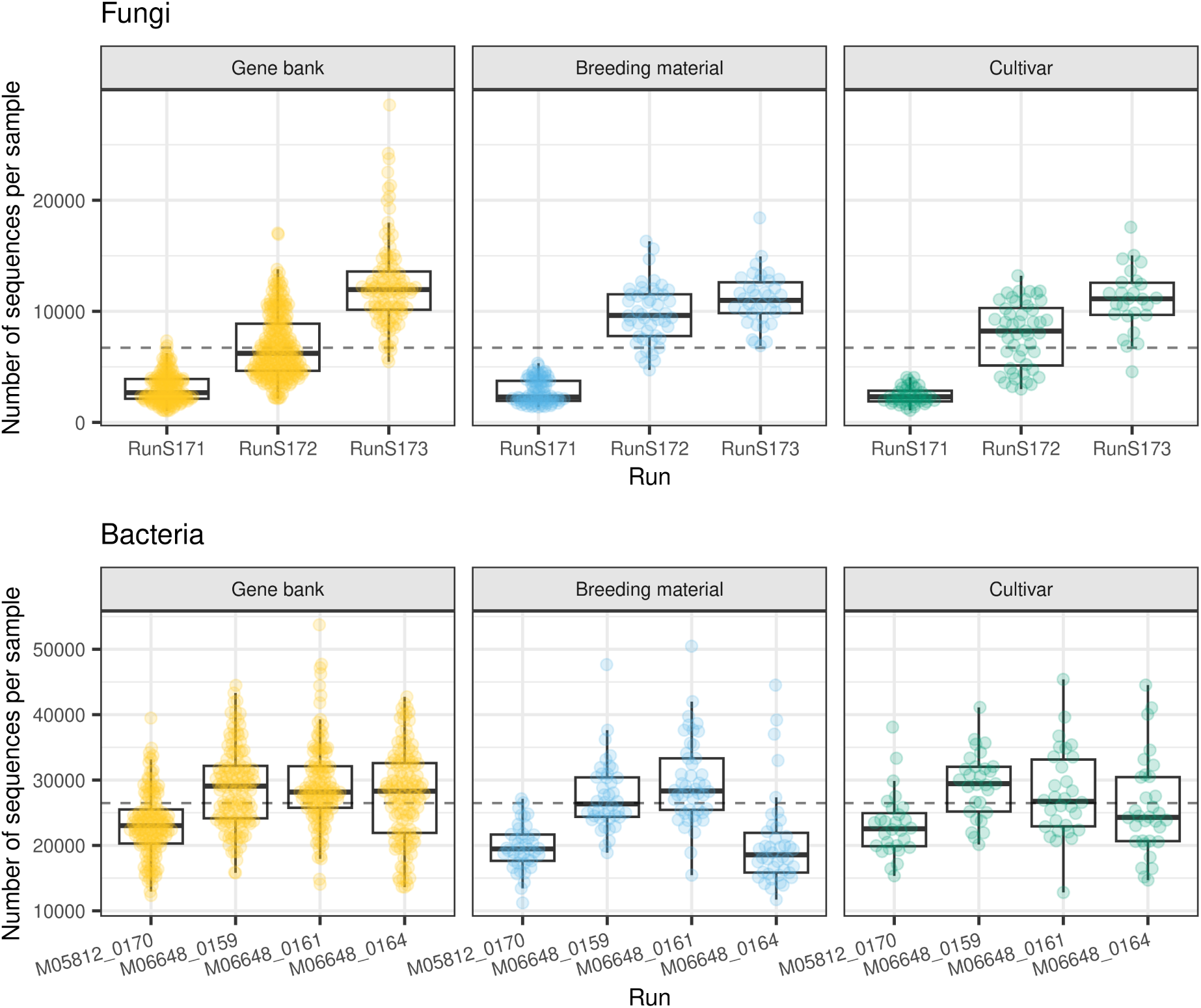
Sequencing depth of amplicon sequencing data for fungi (top panel) and bacteria (bottom panel) grouped by seed source and sequencing run. Dashed line indicates the mean sequencing depth for fungi (top) and bacteria (bottom).

**Figure S3.**
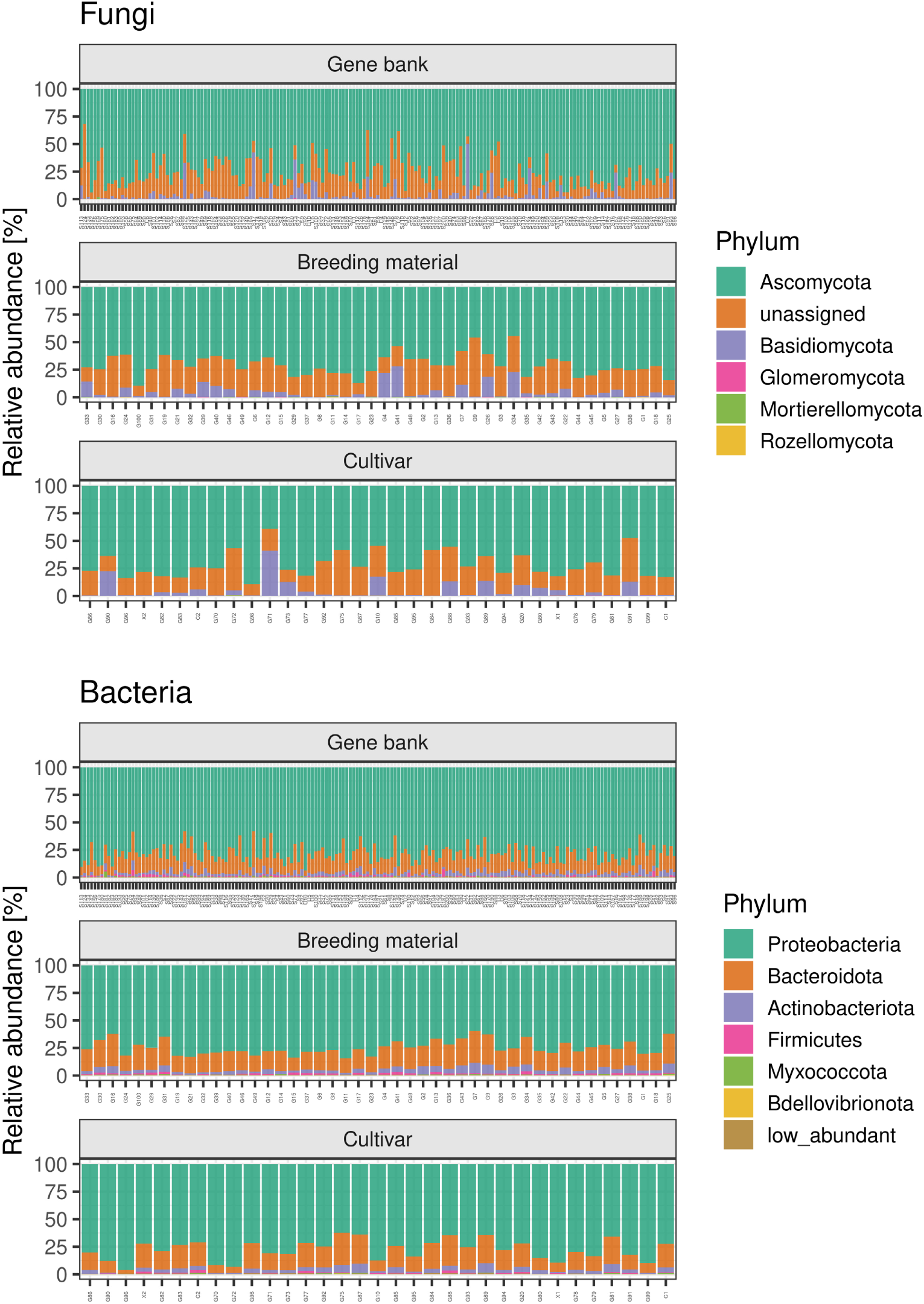
Taxonomy of genotypes at Phylum level for fungi (top panel) and bacteria (bottom panel). Genotypes are grouped by seed source and ordered by increasing emergence level.

**Figure S4.**
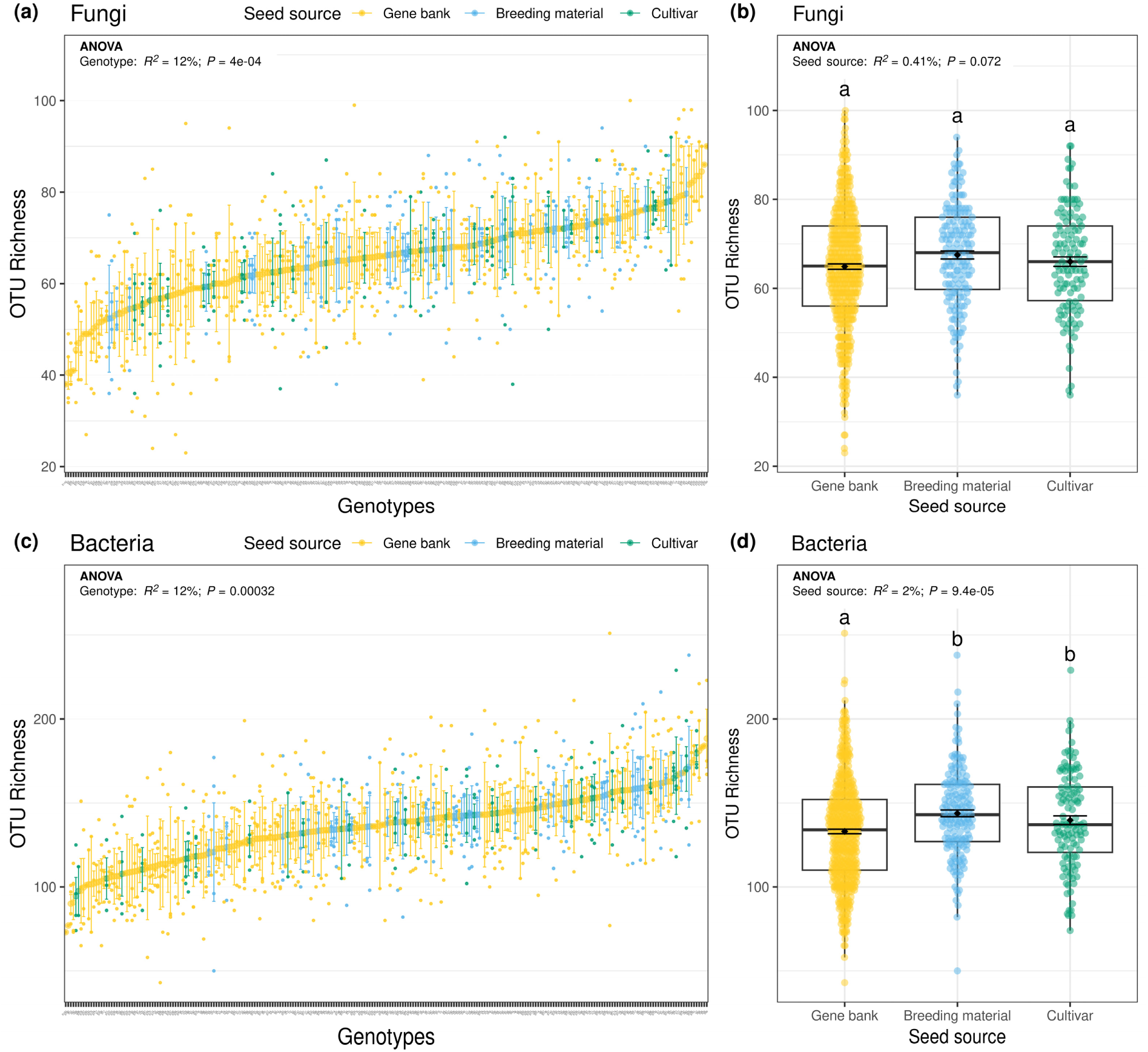
Influence of plant genotype **(a, c)** and seed source **(b, d)** on root fungal **(a, b)** and bacterial (c, d) OTU richness. All plots show individual datapoints, means ±SE, and the ANOVA tables with the explained variance (adjusted R^2^) and the corresponding *P* value. In **(b)** and **(d)** boxplots are also shown. Letters indicate significant differences among seed sources (analysis of variance followed by pairwise comparison of estimated marginal means, *P*_adj_ < 0.05).

**Figure S5.**
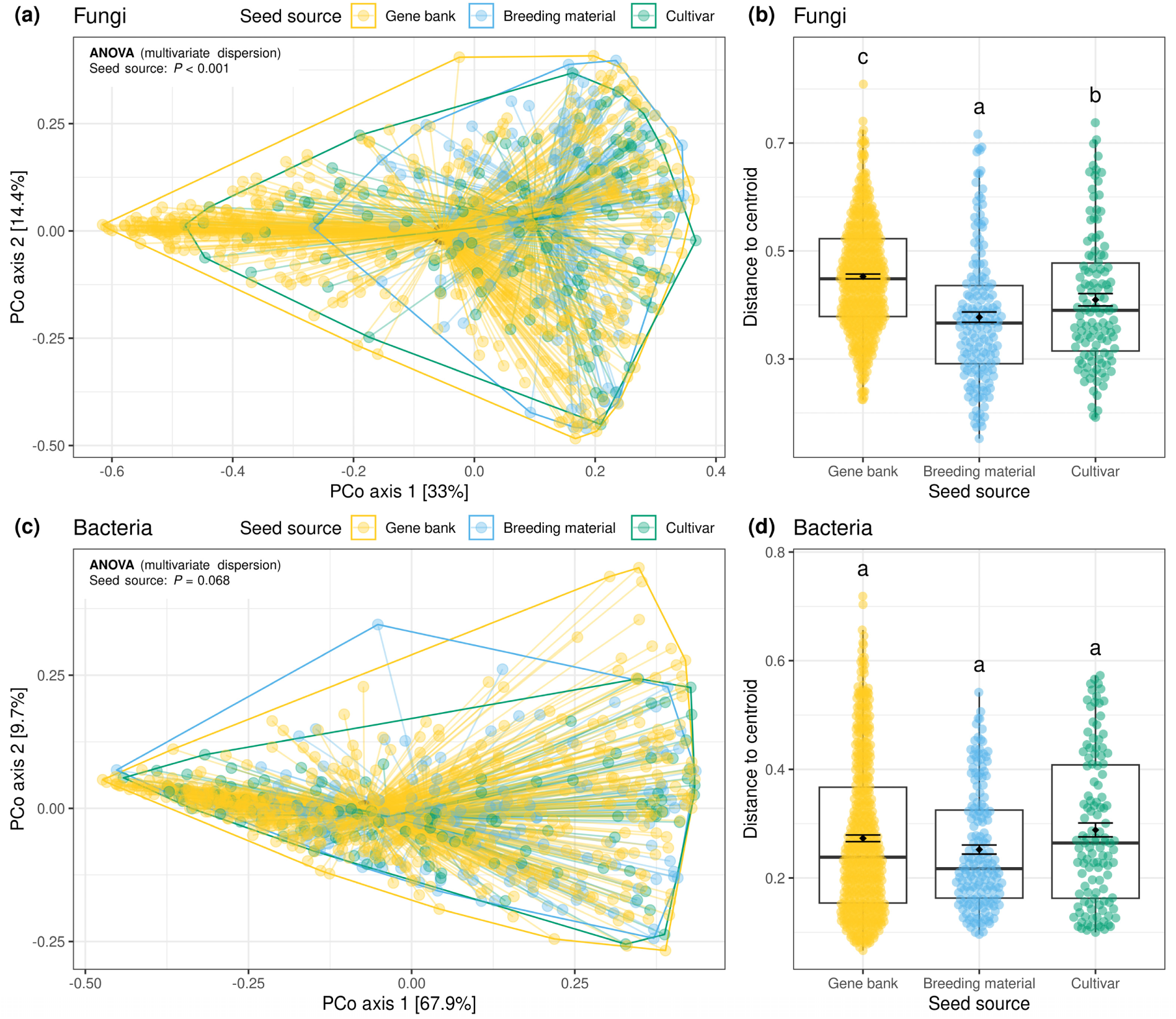
Influence of seed source on root fungal **(a, b)** and bacterial **(c, d)** multivariate dispersion (variance) assessed by **(a)** Principal Coordinates Analysis (PCoA) ordination and **(b)** distances to group centroid. ANOVA results of multivariate dispersion (*betadisper*) with the corresponding *P* value are included in both plots. All plots show individual datapoints, in **(b)** and **(d)** boxplots are also shown. Letters indicate significant differences among seed sources (analysis of variance followed by pairwise comparison of estimated marginal means, *P*_adj_ < 0.05).

**Figure S6.**
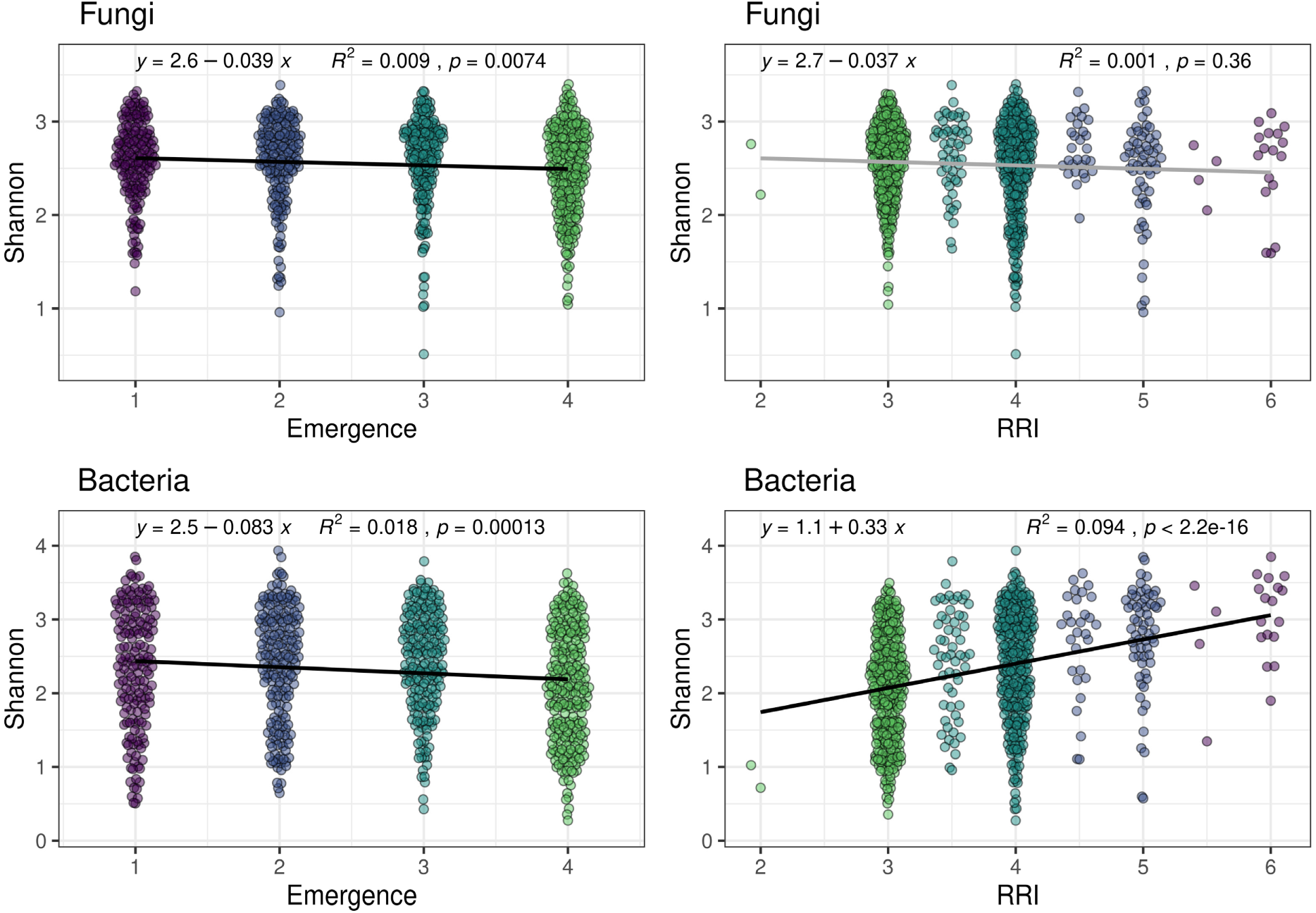
Correlation between root fungal (top) and bacterial (bottom) Shannon diversity and root rot resistance. Correlations are shown for emergence (left panels) and root rot index (RRI, right panel). All plots show individual datapoints, the explained variance (R^2^), the corresponding *P* value derived from a Spearman correlation, and the linear regression equation.

**Figure S7.**
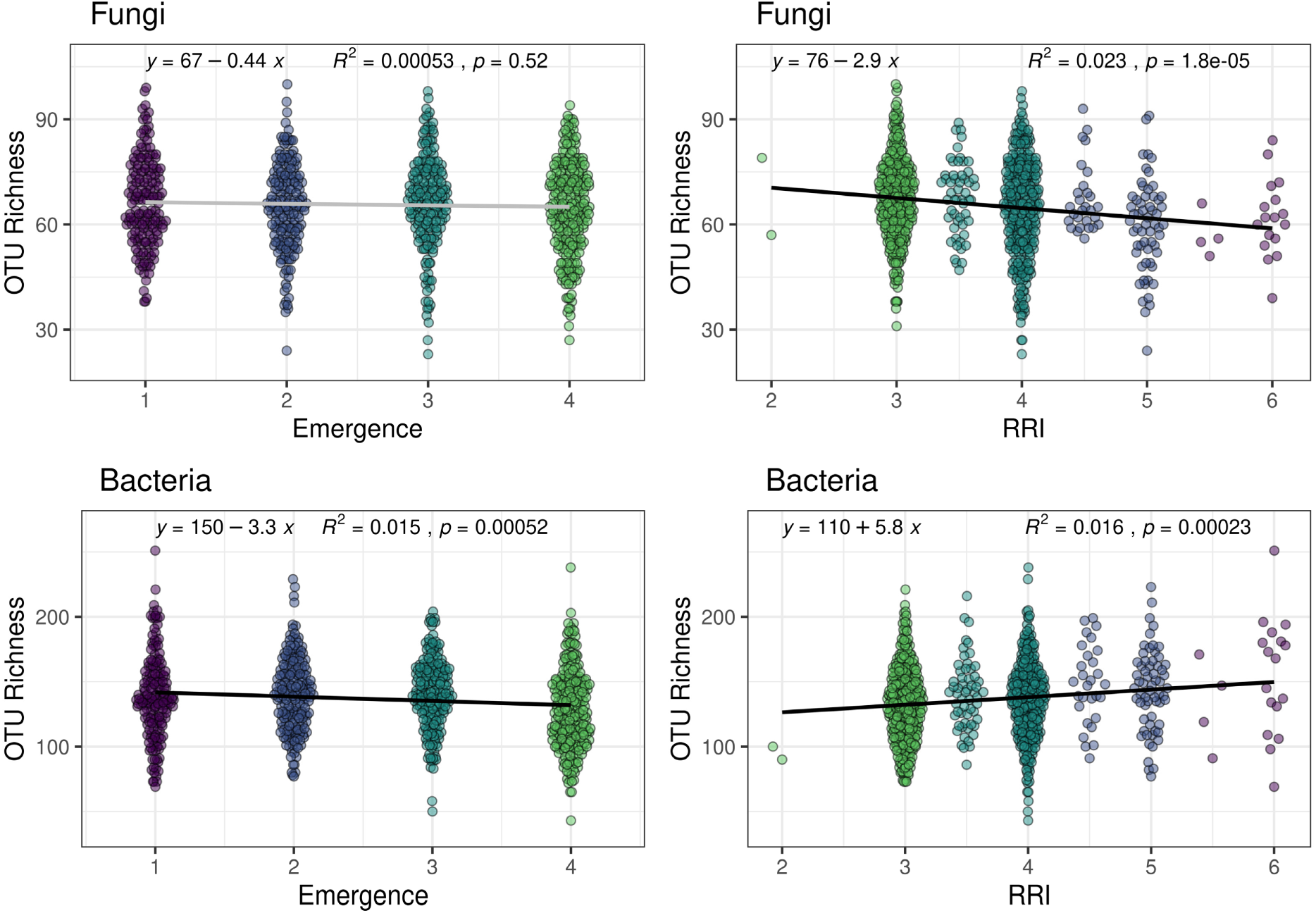
Correlation between root fungal (top) and bacterial (bottom) OTU richness and root rot resistance. Correlations are shown for emergence (left panels) and root rot index (RRI, right panel). All plots show individual datapoints, the explained variance (R^2^), the corresponding *P* value derived from a Spearman correlation, and the linear regression equation.

**Figure S8.**
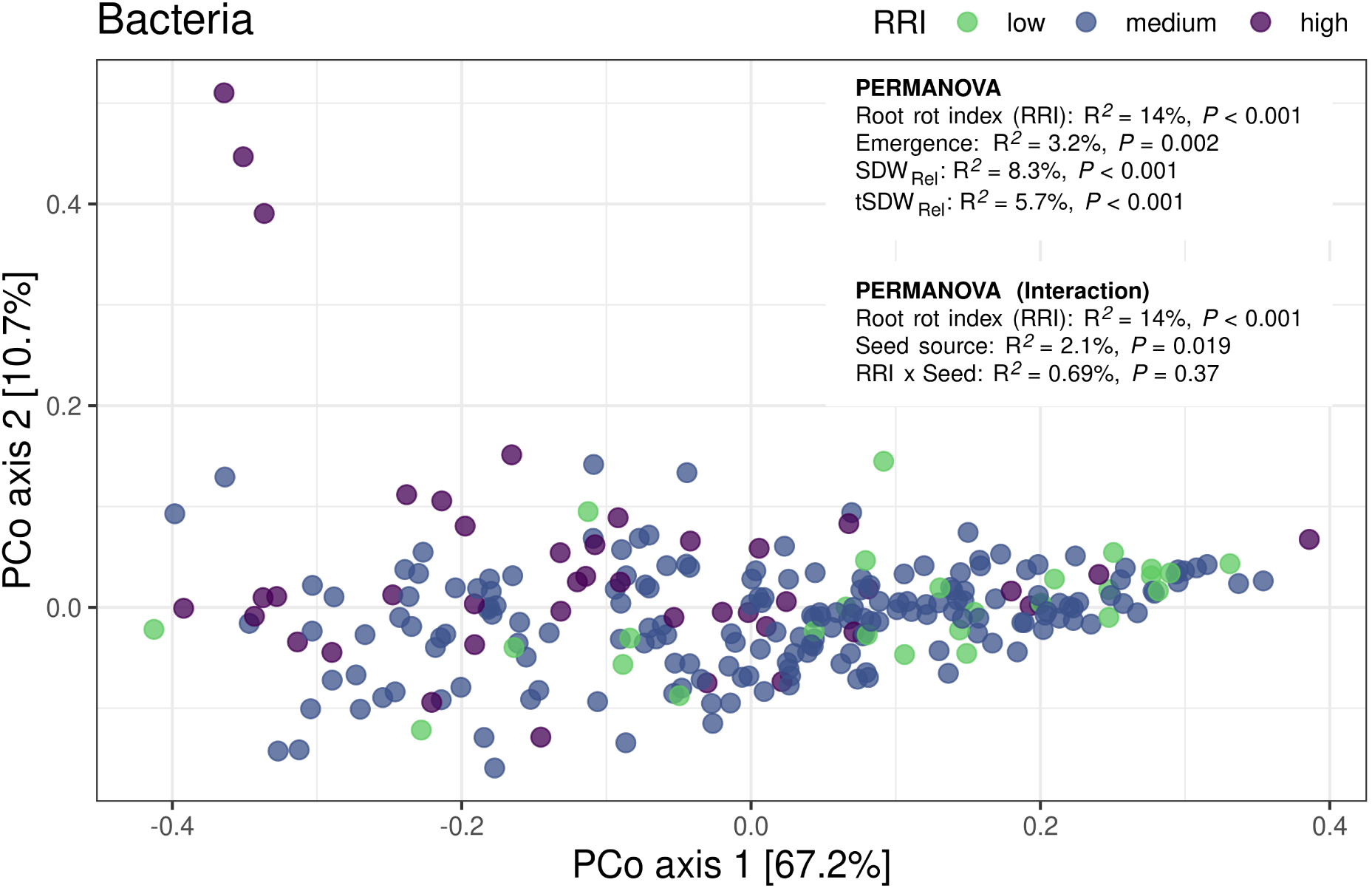
Association of root bacterial beta diversity of genotype means with root rot resistance assessed by Principal Coordinates Analysis (PCoA) ordination. PERMANOVA results with the explained variances (R^2^) and the corresponding *P* values for different root rot resistance-associated traits (top) and the interaction with seed source (bottom) are included. SDW_Rel_: Relative shoot dry weight per plant, tSDW_Rel_: Relative total shoot dry weight per pot.

**Figure S9.**
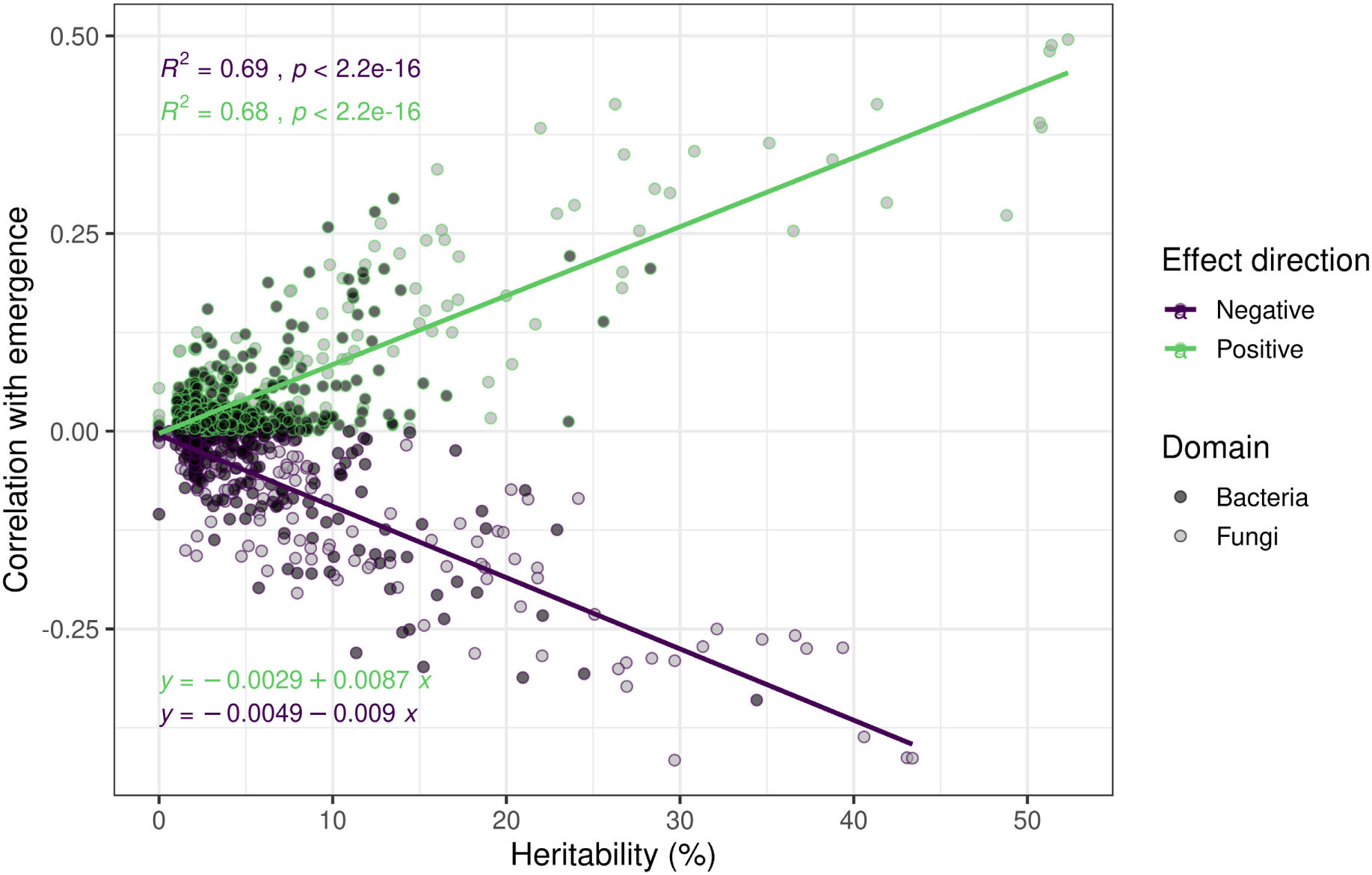
Correlation between heritability and magnitude of correlation with root rot resistance for OTUs with negative (green) or positive (purple) association with emergence. Explained variances (R^2^) and the corresponding *P* values (top) and linear regression equation (bottom) are shown.

**Figure S10.**
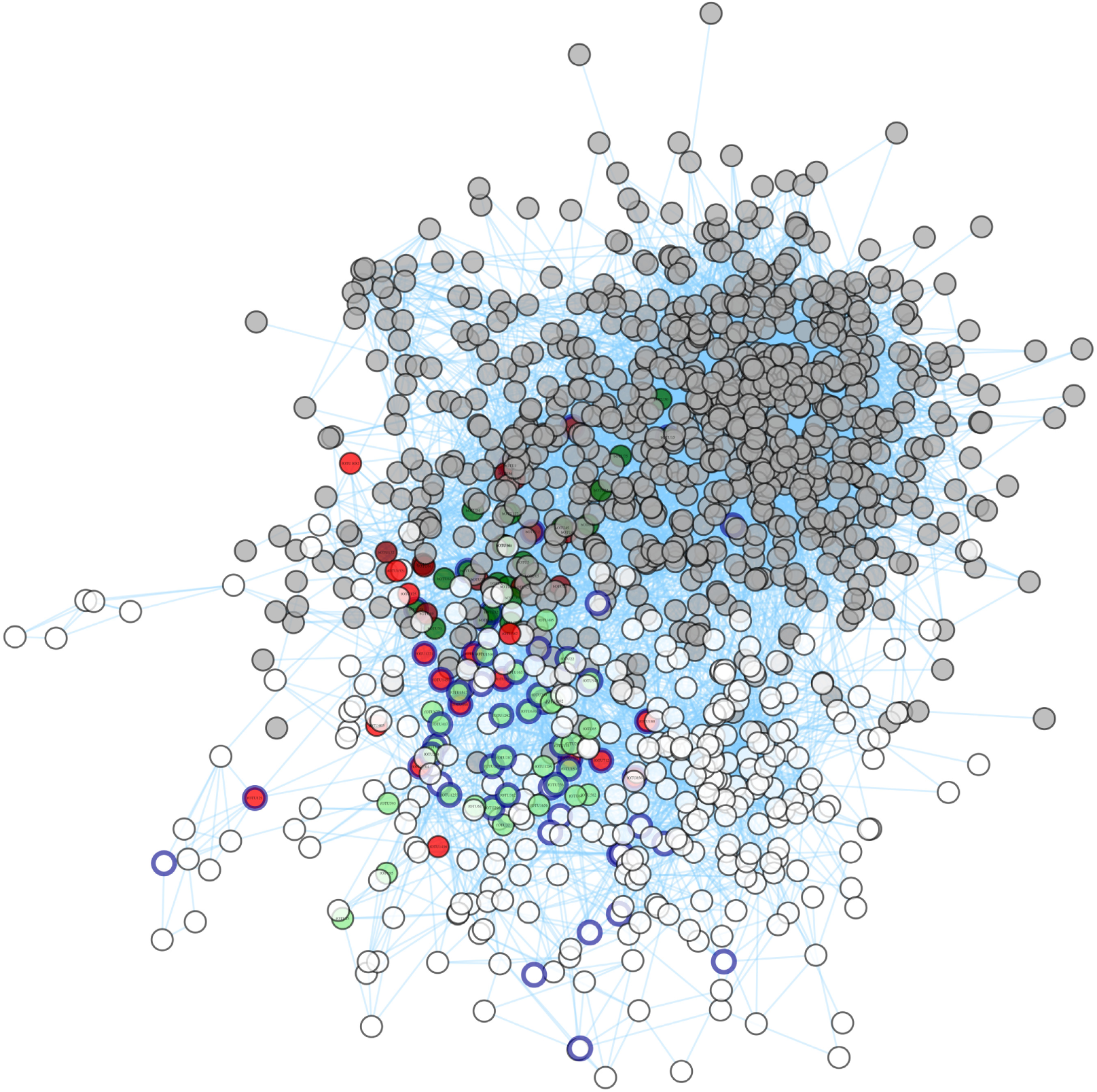
Co-occurrence network of bacterial and fungal OTUs. Gray, dark red and dark green nodes belong to fungal OTUs. White, light red and light green nodes belong to bacterial OTUs. Red nodes are significantly associated with root rot susceptibility, green nodes are significantly associated with root rot resistance. Blue framed nodes represent the top 58 heritable OTUs.

**Figure S11.**
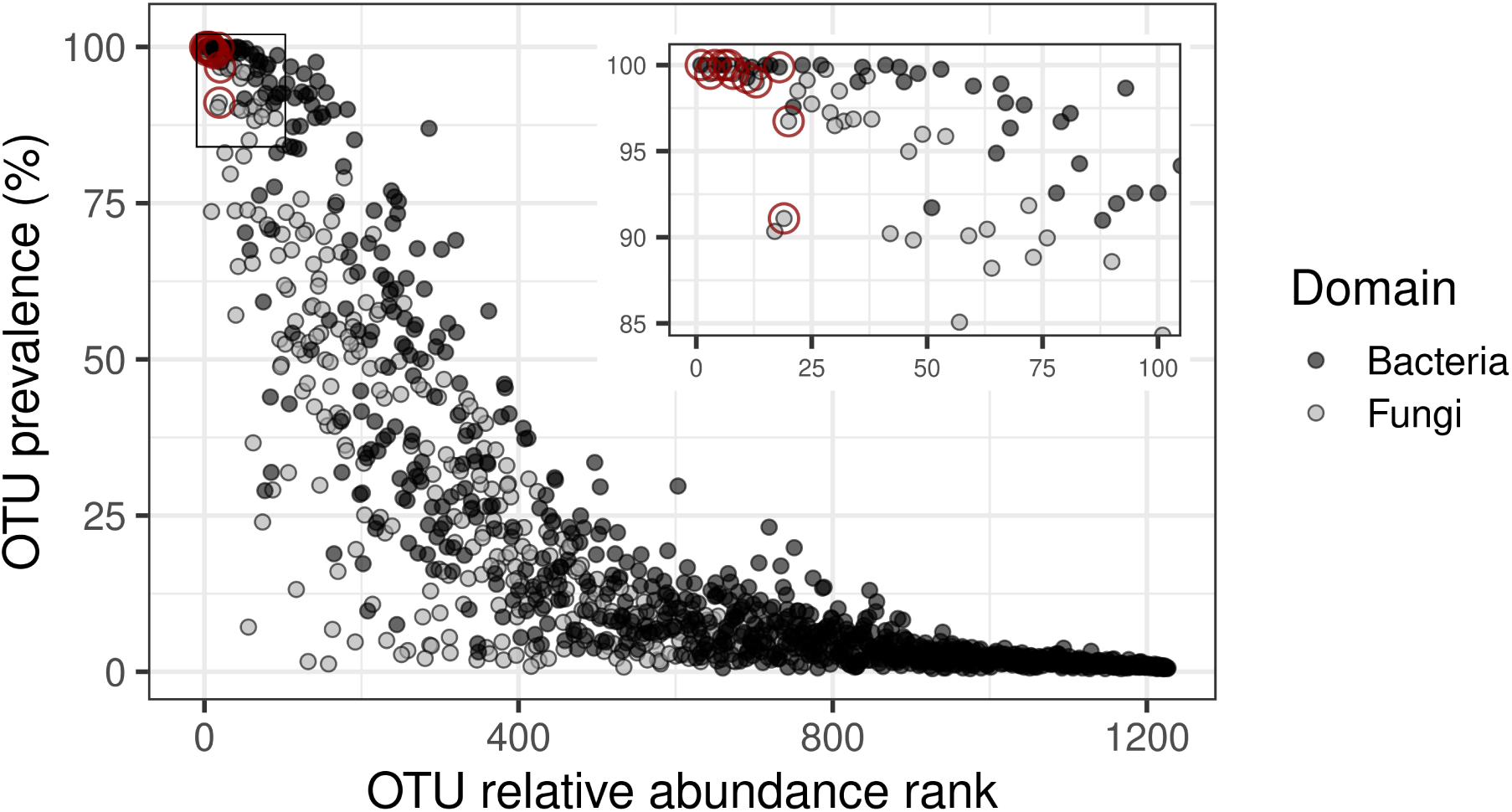
Prevalence plot showing each OTUs relative abundance rank and prevalence. The 11 most resistance-associate, heritable, and connected OTUs (shown in **Figure 5 e**) are highlighted in red.

**Figure S12.**
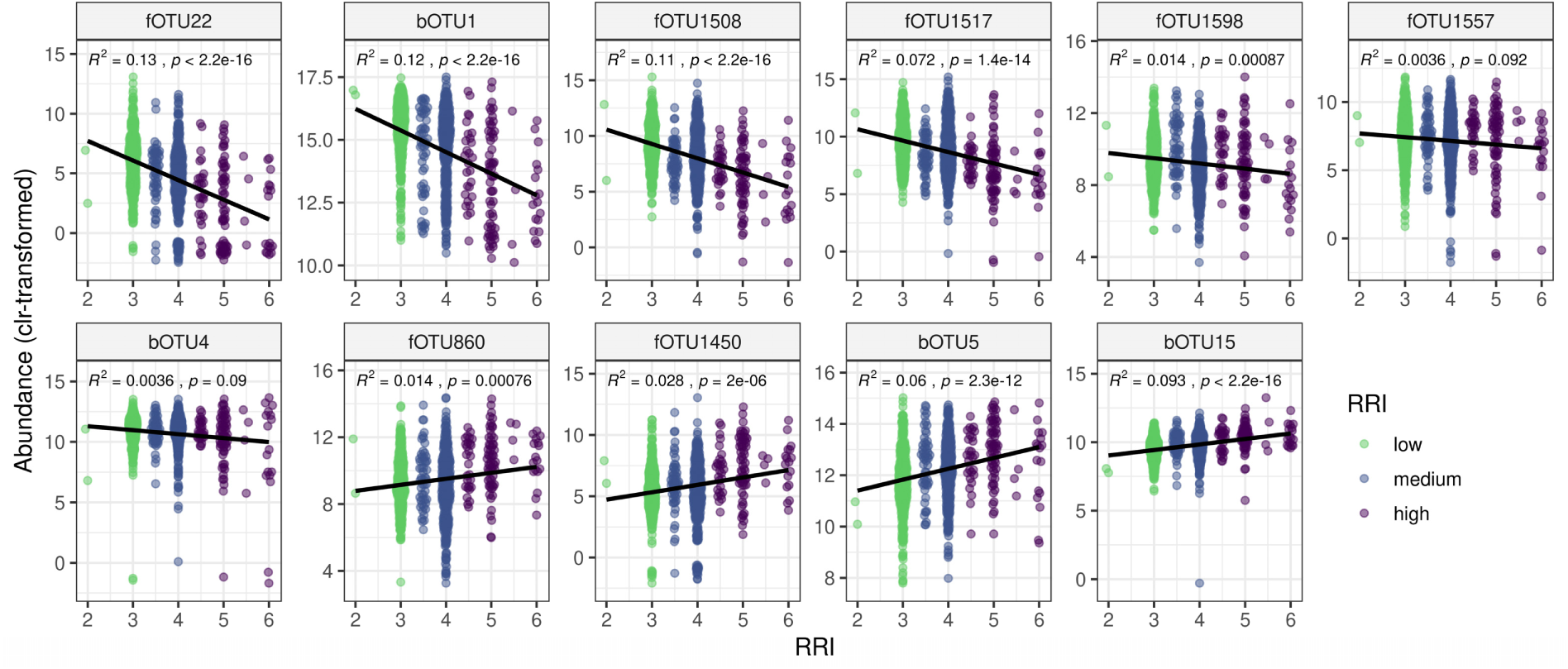
Correlation between root rot index (RRI) and OTU abundance for OTUs with high relative abundance, connectedness in the microbial network, heritability and association with resistance. Explained variance (R^2^) and *P* values of correlations are indicated. SDW_Rel_: Relative shoot dry weight per plant, tSDW_Rel_: Relative total shoot dry weight per pot. clr-transformed: centered log-ratio transformed.

### Supplementary Tables

**Supplementary Table 1.** Attributes of OTUs previously reported to be root rot associated. Taxonomy, relative abundance (RA), heritability, prevalence, network characteristics egree and betweenness) and correlation (Spearman) with root rot resistance-associated traits of OTUs are listed. *P* values < 0.05 are highlighted in bold.

| OTU id | Phylum | Family | Genus | Blast hit<sup>A</sup> | RA (%)<sup>B</sup> | Heritability (%) | Prevalence (%) | Degree | Betweenness | Emergence |  | RRI |  |
| --- | --- | --- | --- | --- | --- | --- | --- | --- | --- | --- | --- | --- | --- |
|  |  |  |  |  |  |  |  |  |  | cor | *P* | cor | *P* |
| fOTU40 | Ascomycota | Didymellaceae | unassigned | *Ascochyta pisi* | 1.05 | 2.03 | 83.1 | 9 | 356 | -0.154 | < **0.001** | -0.026 | 0.639 |
| fOTU419 | Ascomycota | Nectriaceae | Fusarium | *Fusarium sp.* | 0.745 | 15.7 | 73.8 | 8 | 1850 | -0.138 | < **0.001** | 0.016 | 0.775 |
| fOTU831 | Basidiomycota | Ceratobasidiaceae | unassigned | *Rhizoctonia sp.* | 3.86 | 3.02 | 73.7 | 7 | 40.5 | -0.114 | **0.006** | 0.003 | 0.907 |
| fOTU680 | Ascomycota | Nectriaceae | Fusarium | *Fusarium sp.* | 0.095 | 20.7 | 52.7 | 6 | 630 | -0.084 | 0.074 | 0.017 | 0.711 |
| fOTU578 | Basidiomycota | Ceratobasidiaceae | unassigned | *Rhizoctonia sp.* | 0.035 | 5.73 | 22.2 | 10 | 639 | -0.048 | 0.382 | 0.024 | 0.616 |
| fOTU335 | Basidiomycota | Ceratobasidiaceae | unassigned | *Ceratobasidium sp.* | 0.108 | 2.01 | 13.2 | 3 | 358 | -0.023 | 0.574 | 0.027 | 0.557 |
| fOTU329 | Ascomycota | Nectriaceae | Fusarium |  | 0.004 | 2.53 | 3.01 | 17 | 3130 | -0.018 | 0.597 | 0.04 | 0.447 |
| fOTU140 | Basidiomycota | Ceratobasidiaceae | unassigned | *Rhizoctonia sp.* | 0.096 | 2.04 | 1.63 | 5 | 98.3 | 0.002 | 0.611 | 0.044 | 0.43 |
| fOTU334 | Ascomycota | Nectriaceae | Fusarium | *Fusarium venenatum* | 0.005 | 1.97 | 8.41 | 3 | 27.5 | 0.027 | 0.549 | 0.01 | 0.609 |
| fOTU137 | Ascomycota | Nectriaceae | Fusarium | *Fusarium redolens* | 0.142 | 2.01 | 70 | 4 | 116 | 0.03 | 0.581 | -0.094 | **0.05** |
| fOTU487 | Ascomycota | Nectriaceae | Fusarium | *Fusarium sp.* | 0.008 | 3.4 | 19.1 | 5 | 1010 | 0.044 | 0.392 | 0 | 0.681 |
| fOTU1020 | Glomeromycota | Glomeraceae | Funnelformis |  | 0.037 | 20.8 | 45 | 17 | 2590 | 0.084 | 0.1 | -0.236 | < **0.001** |
| fOTU60 | Ascomycota | Bionectriaceae | unassigned | *Clonostachys rosea* | 0.501 | 11.6 | 89.8 | 7 | 688 | 0.1 | **0.018** | -0.082 | 0.086 |

<sup>A</sup>Top hit of a BLAST search against the NCBI database; <sup>B</sup>Within-Kingdom relative abundance was computed for each OTU.

**Supplementary Table 2.** Model comparison between models explaining seedling emergence. Stepwise regression on most abundant OTUs (20 fungi and 20 bacteria) and beta diversity (PCo axis 1 and 2 for fungi and bacteria), most abundant OTUs alone and a simple linear model of the most correlated PCo axis and the most correlated OTU were performed (rows). The number of variables in the model, the Akaike’s information criterion (AIC), the adjusted R^2^ and the *P* value of the model are shown.

| Model (emergence ~) | Nr. of variables | AIC | Adjusted R<sup>2</sup> (%) | *P* value |
| --- | --- | --- | --- | --- |
| Stepwise regression OTUs and PCoA axes<sup>A</sup> | 15 | 2098.7 | 36.7 | 1.61e-70 |
| Stepwise regression OTUs<sup>B</sup> | 12 | 2144.5 | 32.7 | 4.37e-62 |
| PCoA axis 1 of fungi | 1 | 2192.5 | 27.5 | 1.04e-57 |
| fOTU1508 | 1 | 2242.5 | 22.8 | 7.7e-47 |

<sup>A</sup> Variables in final model: bOTU1, fOTU1517, bOTU2, bOTU7, fOTU731, fOTU1450, fOTU174, fOTU340, bOTU16, bOTU9, fOTU856, bOTU18, bPCoA\_2, fPCoA\_1, fPCoA\_2
<sup>B</sup> Variables in final model: fOTU1508, fOTU1063, bOTU2, bOTU7, fOTU731, fOTU340, bOTU11, bOTU9, bOTU18, bOTU33, fOTU1092, fOTU712

**Supplementary Table 3.** Model comparison between models explaining root rot index (RRI). Stepwise regression on most abundant OTUs (20 fungi and 20 bacteria) and beta diversity (PCo axis 1 and 2 for fungi and bacteria), most abundant OTUs alone and a simple linear model of the most correlated PCo axis and the most correlated OTU were performed (rows). The number of variables in the model, the Akaike information criterion (AIC), the adjusted R^2^ and the *P* value of the model are shown.

| Model (RRI ~) | Nr of variables | AIC | Adjusted R<sup>2</sup> (%) | *P* value |
| --- | --- | --- | --- | --- |
| Stepwise regression OTUs and PCoA axes<sup>A</sup> | 19 | 1441.8 | 33.9 | 2.41e-61 |
| Stepwise regression OTUs<sup>B</sup> | 19 | 1441.8 | 33.9 | 2.41e-61 |
| PCoA axis 1 of bacteria | 1 | 1664.5 | 10.6 | 3.07e-21 |
| bOTU1 | 1 | 1623.6 | 15 | 3.47e-30 |

<sup>A</sup> Variables in final model: bOTU1, fOTU1508, fOTU1517, bOTU5, fOTU1598, bOTU15, fOTU731, fOTU1450, fOTU174, bOTU6, bOTU21, fOTU1557, bOTU16, bOTU9, bOTU13, fOTU630, fOTU1092, fOTU1556, bOTU12
<sup>B</sup> Variables in final model: bOTU1, fOTU1508, fOTU1517, bOTU5, fOTU1598, bOTU15, fOTU731, fOTU1450, fOTU174, bOTU6, bOTU21, fOTU1557, bOTU16, bOTU9, bOTU13, fOTU630, fOTU1092, fOTU1556, bOTU12

